# Colony stimulating factor 1 signaling regulates myeloid fates in zebrafish via distinct action of its receptors and ligands

**DOI:** 10.1101/2021.04.06.438628

**Authors:** Martina Hason, Tereza Mikulasova, Olga Machonova, Antonio Pombinho, Tjakko J van Ham, Uwe Irion, Christiane Nüsslein-Volhard, Petr Bartunek, Ondrej Svoboda

## Abstract

Macrophage colony-stimulating factor receptor (M-CSFR/CSF1R) signaling is crucial for the differentiation, proliferation, and survival of myeloid cells. Therapeutic targeting of the CSF1R pathway is a promising strategy in many human diseases, including neurological disorders or cancer. Zebrafish are commonly used for human disease modeling and preclinical therapeutic screening. Therefore, it is necessary to understand the proper function of cytokine signaling in zebrafish to reliably model human-related diseases. Here, we investigate the roles of zebrafish Csf1rs and their ligands - Csf1a, Csf1b and Il34, in embryonic and adult myelopoiesis. The proliferative effect of exogenous Csf1a on embryonic macrophages is connected to both receptors as it is diminished in both *csf1ra*^Δ5bp^ and *csf1rb*Δ^4bp^ mutants. There is no evident effect of Csf1b in zebrafish embryonic myelopoiesis. Further, we uncover an unknown role of Csf1rb in zebrafish granulopoiesis. Deregulation of Csf1rb signaling leads to failure in myeloid differentiation resulting in neutropenia throughout the whole lifespan. Surprisingly, Il34 signaling through Csf1rb seems to be of high importance as both *csf1rb*^Δ4bp^ and *il34*^Δ5bp^ deficient zebrafish larvae lack granulocytes. Our single-cell RNA sequencing analysis of adult whole kidney marrow (WKM) hematopoietic cells suggests that *csf1rb* is expressed mainly by blood and myeloid progenitors and that the expression of *csf1ra* and *csf1rb* is non-overlapping. We point out differentially expressed genes important in hematopoietic cell differentiation and immune response in selected WKM populations. Our findings could improve the understanding of myeloid cell function and lead to the further study of CSF1R pathway deregulation in disease, mostly in cancerogenesis.

**Key points:**

- *csf1ra* and *csf1rb* are indispensable for macrophage differentiation and together with *csf1a* regulate embryonic macrophage fates in zebrafish
- *csf1rb* is important for granulocyte differentiation and migration and together with *il34* it regulates embryonic granulocytic fates in zebrafish

## Introduction

Hematopoiesis is a process of proliferation, differentiation, fate-commitment, and self-renewal of blood cells. It is primarily regulated by extrinsic signals, such as cytokines and growth factors that bind to cell receptors and activate internal signaling pathways.^1,2^ One of the most prominent receptors that control the myeloid compartment is colony-stimulating factor 1 receptor (CSF1R, also known as macrophage-CSFR). In mammals, it is activated upon binding of two distinct cytokine ligands that have no obvious sequence homology – colony stimulating factor 1 (CSF1, M-CSF) and interleukin 34 (IL-34).^3–6^ However, despite the fact that both of these cytokines bind to the same receptor and can equally support cell growth and survival, they achieve this by triggering different chemokine responses.^4,7^ The CSF1R signaling pathway is in general critical for the proliferation, differentiation, survival, and activation of mononuclear phagocytic cells (MPCs) such as monocytes, macrophages, osteoclasts, or microglia in mammals,^8–10^ birds,^11,12^ and fish.^13^ Deregulation of the CSF1R pathway was connected to disease phenotypes (reviewed in Hume et al.^14^) such as osteopetrosis,^9,10,15,16^ brain disease,^17–20^ or cancer.^21–24^ Thus the CSF1R signaling is of high interest as a pathway for therapeutic targeting in neurological and infectious diseases and in tumorigenesis.^25,26^ Particularly, myeloid cells, including both neutrophils and macrophages, can act negatively in carcinogenesis. Therefore, tumor associated macrophages (TAMs) are believed to be critical in tumor metastasis and are a good target in addition to conventional chemotherapy.^27–29^ It has been shown in mouse that the number of TAMs can be efficiently reduced by the inhibition of CSF1R. Because of its low-throughput when testing compounds, other model organisms need to be utilized.^30^ Zebrafish is a convenient model organism for human disease modeling^31–33^ and the small size of zebrafish makes it advantageous for high-throughput pre-clinical drug screening.^32,34–38^ Due to the genome re-duplication in teleost fish, many paralogs were generated that could possess redundant or novel biological functions.^39–41^ This includes both, Csf1 and Csf1r, and therefore it is still needed to define the role and specificity of Csf1a, Csf1b and Il34 towards Csf1rs (Csf1ra/b) in zebrafish myelopoiesis. So far, it seems that the function of Csf1ra and Csf1rb is only partially redundant.^15^ For instance, there are spatiotemporal differences in the importance of Csf1rs for microglia and HSC-derived myeloid cells development and seeding of the zebrafish brain.^42–44^

In this article, we focus on the roles of Csf1a, Csf1b, and Il34 cytokines in zebrafish embryonic myelopoiesis and adult hematopoiesis, shown by *ex vivo* tools and single cell RNA sequencing (scRNA-seq) of whole kidney marrows (WKM). We use a collection of zebrafish loss-of-function mutants to discern the effects of Csf1-receptor and -ligand functional defects. We show that Csf1a drives the expansion of embryonic macrophages, Csf1b has no evident role in embryonic myelopoiesis and Il34, acting through Csf1rb, is important for embryonic granulopoiesis. Finally, our observations suggest evolutionarily interesting functions of CSF1R signaling in the myelopoiesis of non-mammalian vertebrates in addition to the conventional role of CSF1 in mammalian myelopoiesis^8,9,45^ that should be taken into consideration when modelling human myeloid disorders in zebrafish.

## Materials and methods

### Animals

Zebrafish were bred, raised and kept in ZebTEC aquatic systems (Tecniplast) according to standard procedures,^46^ tracked using Zebrabase.^47^ Zebrafish *csf1* receptor mutant lines used in this study were *csf1ra*^V614M^ (Panther),^48^ *csf1ra*^t36ui^, (further *csf1ra*^Δ5bp^), *csf1rb*^re01^, (further *csf1rb*^Δ4bp^)^42^ and *csf1ra*^V614M^;*csf1rb*^re01^ double mutants.^42^ The *csf1r* ligand mutants used were *csf1a*^ins2bp^, *csf1b*^Δ2bp^, and *il34*^Δ5bp^.^49^ Transgenic reporter zebrafish lines used were Tg(*mpeg1:EGFP*),^50^ Tg(*fms:GAL4;UAS:mCherry*),^51^ Tg(*mpx:EGFP*),^52^ and Tg(*pax7:GFP*).^53^ WT(AB) were used as controls. For *ex vivo* experiments, 6–12-month-old fish were used to get an optimal number of whole kidney marrow (WKM) cells. Animal care and experiments were approved by the Animal Care Committee of the Institute of Molecular Genetics, Czech Academy of Sciences (13/2016 and 96/2018) in compliance with national and institutional guidelines.

### Multiplexed quantitative RNA fluorescence *in situ* hybridization

Hybridization chain reaction (HCR) v3.0 probe sets, amplifiers, and buffers, were used according to the manufacturer’s protocols (Molecular Instruments).^54^ Probes detecting zebrafish *csf1rb* (XM_009295703.3), *mpeg1* (NM_212737.1) and *mpx* (NM_001351837.1) were designed by the manufacturer. The Alexa 647, Alexa 546 and Alexa 488 amplifiers were used.

### Fluorescence imaging

Fluorescent images were acquired on Zeiss Axio Zoom.V16 with Axiocam 506 mono camera. Orthogonal projections were created in ZEN Blue 2.3 software. Images of HCR-stained embryos were acquired on, Dragonfly 503 microscope (Andor) using Zyla 4.2 PLUS sCMOS camera. All images were processed by Fiji and Adobe Photoshop CC 2021.^55^

### Single cell RNA sequencing (scRNA-seq) and transcriptomics

WKM cells were isolated as described previously ^56^, fractionated with Biocoll (1.077 g/ml, Merck) and counted. Between 3,000 and 5,000 cells were used for preparation of Chromium 3’ sequencing libraries using Chromium Single Cell 3’ Chip kit v3.1 and sequenced with Illumina Nextseq 500. The Illumina FASTQ files were used to generate filtered matrices using CellRanger (10X Genomics) with default parameters. To generate filtered matrices, data were loaded to Cellbender package^57^ using the following parameters - expected-cells = 5000, total-droplets-included = 15000. Filtered matrices were then imported into R for exploration and statistical analysis using a Seurat V3 package.^58^ Counts were normalized according to total expression, multiplied by a scale factor (10,000), and log-transformed. For cell cluster identification and visualization, gene expression values were also scaled according to highly variable genes after controlling for unwanted variation generated by sample identity. Cell clusters were identified based on UMAP of the first 20 principal components of PCA using Seurat’s method, FindClusters, with an original Louvain algorithm and resolution parameter value 0.5. Following quality control and basic clustering of each sample, we subsetted individual datasets to contain 1,700 cells each and merged them together. To visualize marker gene expression, Seurat’s method, Dot-Plot, was used. To merge individual datasets and to remove batch effects, Seurat V3 Integration and Label Transfer standard workflow were used.^58^

### Other procedures and methods

The description of mRNA and protein microinjections, whole mount in situ hybridization (ISH) of zebrafish embryos using digoxigenin-labeled antisense riboprobes, cloning of constructs for recombinant protein expression, *ex vivo* WKM cell cultures, Sudan Black B (SBB) staining of embryos, fluorescence activated cell sorting (FACS) analysis, image processing, statistical analysis and data sharing statement of presented data are outlined in the supplemental Materials and Methods available in the online version of this article.

### Data sharing statement

sc-RNAseq data are available in ArrayExpress under accession number E-MTAB-10360. Plasmids for cytokine expression are available via Addgene: 168103 (pAc-His-zfCsf1a), 168104 (pAc-His-zfCsf1b), 168105 (pAc-His-zfIl34) and for mRNA expression: 168110 (pCS2-zfCsf1a), 168111 (pCS2-zfCsf1b) and 168112 (pCS2-zfIl34).

## Results

### Zebrafish *csf1ra* and *csf1rb* are expressed from early embryonic development and have distinct expression patterns in adults

To determine whether the spatiotemporal expression pattern of zebrafish *csf1ra* and *csf1rb* overlap in embryonic development, we crossed *fms:GAL4;UAS:mCherry* (simplified as *csf1ra:mCherry*) and *mpeg1:EGFP* reporter lines to generate triple transgenic animals (Figure 1Aa). At 72 hours post fertilization (hpf), almost all *mpeg1:EGFP+* macrophages are also *csf1ra:mCherry+* (Figure 1Ab, 1Ac) and single *csf1ra:mCherry+* cells in the skin are xanthophores,^48^ zebrafish pigment cells.^53,59^ On the contrary, by performing double fluorescent HCR using probes for *mpeg1 and csf1rb* at 72 hpf (Figure 1Ba), we showed that the majority of macrophages are *mpeg1* single-positive with only few *mpeg1* and *csf1rb* double-positive cells (Figure 1Bb-c).

**Figure 1:**
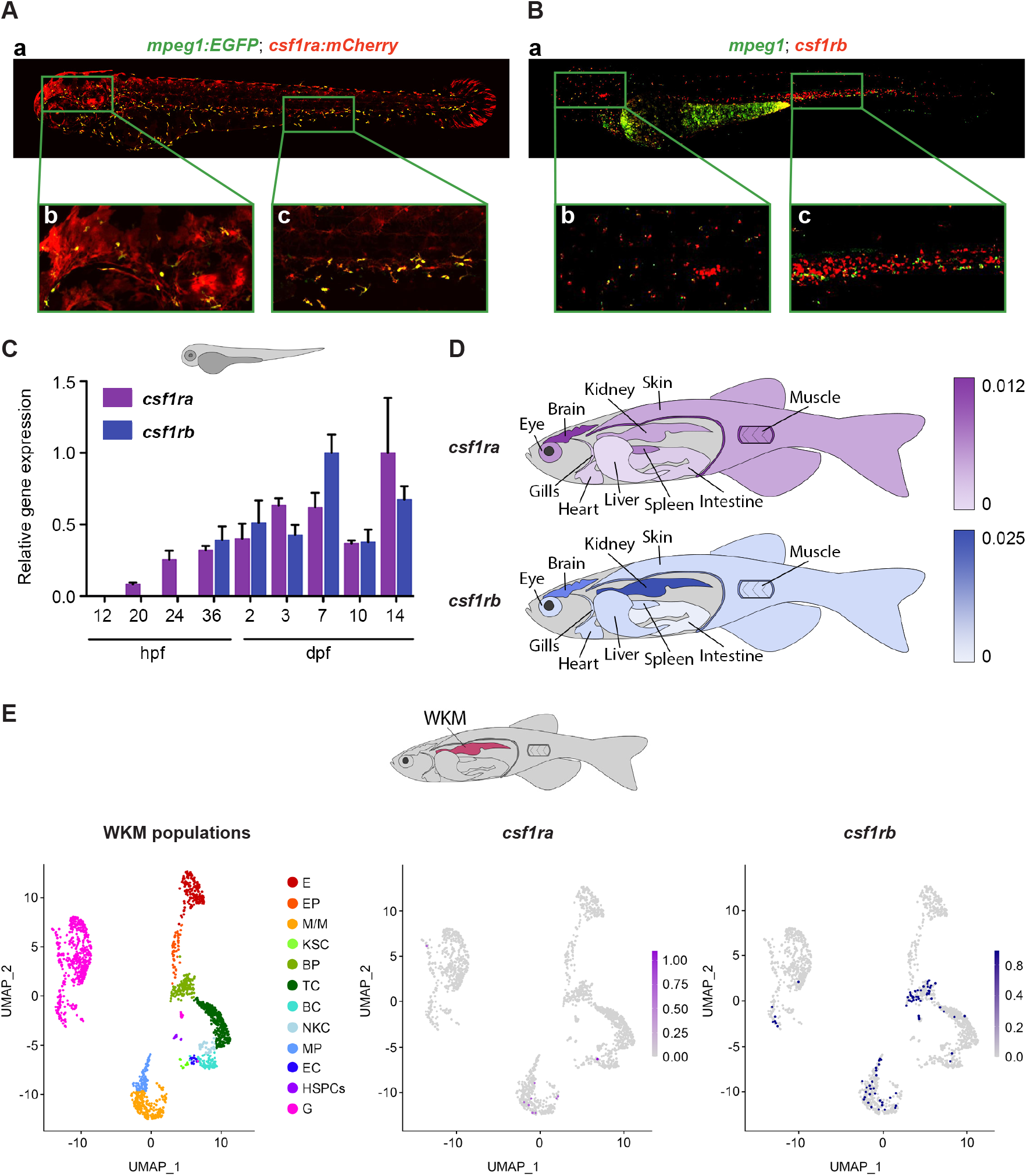
*csf1ra* and *csf1rb* have distinct expression patterns in zebrafish. (A) Co-expression of *csf1ra* (red) and *mpeg1* (green) visualized in 72 hpf Tg(*fms:GAL4;UAS:mCherry*);Tg(*mpeg1:EGFP*) double transgenic embryos: (Aa) whole embryo, (Ab) head, (Ac) caudal hematopoietic tissue (CHT) region. (B) HCR WISH of 72 hpf embryos for *csf1rb* (red) and *mpeg1* (green). (Ba) whole embryo, (Bb) head, (Bc) CHT region. Fluorescence images were taken on Dragonfly 503 microscope (Andor) using Zyla 4.2 PLUS sCMOS camera, with 10x magnification and processed with the Fusion software, FIJI and Adobe Photoshop. (C) qRT-PCR analysis of pooled zebrafish embryos showing the expression dynamics of *csf1ra* and *csf1rb* in zebrafish development. Pool of 15-20 embryos/sample in 2-6 biological replicates. The expression was normalized to *mob4* gene and to the time point with the highest expression (14 dpf for *csf1ra* and 7 dpf for *csfrb*). (D) qRT-PCR analysis of adult zebrafish tissues. Pool of 3-5 fish organs/sample in 3-5 biological replicates. The expression was normalized to *ef1α* gene. (E) scRNA-seq data showing the expression of *csf1ra* and *csf1rb* in whole kidney marrow (WKM) cell populations. E - erythroid cells; EP - erythroid progenitors; M/M - monocytes & macrophages; KSC - kidney support cells; BP - blood progenitors; TC - T-cells; BC - B-cells; NKC - NK cells; MP - myeloid progenitors; EC - endothelial cells; HSPCs - hematopoietic stem and progenitor cells; G – granulocytes.

To characterize the expression pattern of *csf1ra/b* during development and in adult tissues, we performed qRT-PCR. Here, we demonstrate that *csf1ra* starts to be expressed at 20 hpf, whereas *csf1rb* first appears at 36 hpf (Figure 1C). From this time point, the overall expression of both receptors during embryonic development gradually increases until 7 days post fertilization (dpf).

Similarly, qRT-PCR using selected adult zebrafish tissues (Supplemental Figure S1A) showed high expression of *csf1ra* in brain, moderate expression in spleen, muscles, eyes, kidneys and skin and weak expression in the remaining organs. The strongest expression of *csf1rb* was in adult kidney marrow and brain, whereas it was low in other organs. To summarize these results, we created representative schemas (Figure 1D).

To get a more detailed insight into the expression of *csf1ra*/*b* in adult hematopoietic tissues, we performed scRNA-seq (Figure 1E) and demonstrated that there is no overlap between *csf1ra and csf1rb* expression in adult WKM. Instead, the expression of *csf1ra* is restricted to a few cells within the population of monocytes-macrophages, whereas the *csf1rb*+ cells comprise blood and myeloid progenitors, monocytes-macrophages and granulocytes. This is in agreement with our HCR expression data (Figure 1Ba-c), where we show that only a subset of *csf1rb+* cells are macrophages.

### *csf1a* drives the expansion and differentiation of zebrafish embryonic macrophages

To further characterize the effects of *csf1* ligands on hematopoietic cells, we *in vitro* transcribed and injected mRNA for *csf1a*, *csf1b* and *il34* ligands, into 1-cell stage zebrafish embryos and examined their caudal hematopoietic tissue (CHT) region at 72 hpf. We noticed that the overexpression of *csf1a* but not of *csf1b* or *il34* caused expansion of *csf1ra:mCherry+* (Figure 2A) and *mpeg1:EGFP+* macrophages (Figure 2B). These injected embryos had high expression of both *csf1rs* and macrophage specific markers, such as *mpeg1*, *mfap4* and *lcp1* (Supplemental Figure S2A). Increased expression of *lcp1* was also documented by ISH staining using *lcp1* probe (Supplemental Figure S2B). We also noticed that the overexpression of both *csf1a/b* highly increased the number of *csf1ra:mCherry+* cells across the whole fish. We saw the same expansion in the xanthophore-specific *pax7:EGFP* transgenic line (data not shown).

**Figure 2:**
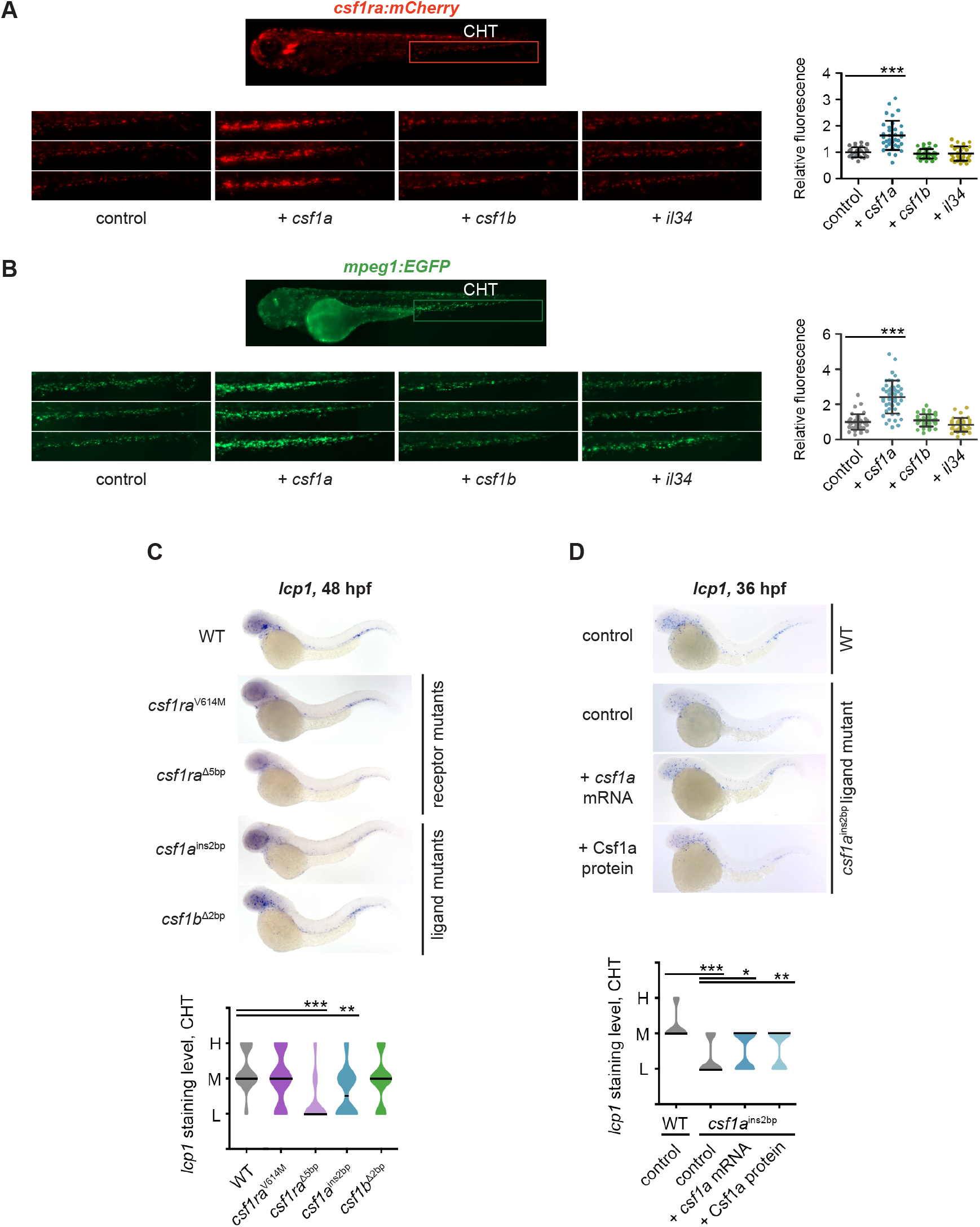
*csf1a,* but not *csf1b* and *il34* drive the expansion of embryonic macrophages *in vivo.* (A-B) *csf1a, csf1b* and *il34* ligands were overexpressed by mRNA microinjection in 1-cell stage transgenic embryos. Control embryos were injected with PBS. Fluorescence images were acquired at 72 hpf and the area of fluorescent cells was calculated in the caudal hematopoietic tissue (CHT, area inside of the red box) by FIJI. Results were normalized to injected controls. Scatter plots on the right represent quantification of fluorescent cells in CHT. Each dot in the scatter plot represents one larva. (A) Tg(*fms:GAL4;UAS:mCherry*) (B) Tg(*mpeg1:EGFP*) (C) WISH of 48 hpf embryos showing the expression of *lcp1* in WT, two *csf1ra* mutants: *csf1ra^V614M^* (panther) and *csf1ra^Δ5bp^*, and in *csf1a^ins2bp^*, *csf1b^Δ2bp^* ligand mutants. (D) *lcp1* WISH of 36 hpf WT embryos injected with PBS (control), and *csf1a^ins2bp^* mutant embryos injected with PBS (control), *csf1a* mRNA or recombinant zebrafish Csf1a protein. (C-D) Violin plot graphs show the level of *lcp1* expression in the CHT region of individual embryos (L = low, M = medium, H = high) with median represented by a black line. (A-D) The level of statistical significance was determined by unpaired two-tailed t-test. *P < 0.04, **P < 0.006, ***P < 0.0001. All fluorescent images were acquired on Zeiss Axio Zoom.V16 with Zeiss Axiocam 506 mono camera and ZEN Blue software. Bright field images of WISH were acquired on Zeiss Axio Zoom.V16 with Zeiss Axiocam 105 color camera and processed using the Extended Depth of Focus module in the ZEN Blue software. FIJI and Adobe Photoshop were used for image processing.

### Embryonic macrophage fate is impaired with the loss of *csf1a* signaling in zebrafish

To study impaired macrophage development upon loss of *csf1ra* or *csf1rb*, we compared *lcp1* expression by ISH in CHT at 48 hpf between WT and other receptor mutants (*csf1ra*^V614M^, *csf1ra*^Δ5bp^, or *csf1rb*^Δ4bp^). While there was no difference in the number of *lcp1* positive cells in the *csf1ra*^V614M^ (Figure 2C), *csf1rb*^Δ4bp^ or in the *csf1ra*^V614M^; *csf1rb*^Δ4bp^ double mutant fish (Supplemental Figure S2C), it was significantly decreased in *csf1ra*^Δ5bp^ mutant animals (Figure 2C).

Even though the number of *lcp1+* cells was unchanged in *csf1rb*^Δ4bp^ mutants, positive cells aggregated more to the rostral part of the CHT as compared to the WT. In addition, we also examined *mpeg1* expression in *csf1ra*^V614M^ as well as in *csf1rb*^Δ4bp^ mutants at 48hpf. As expected, based on published data^43,44^ and *lcp1* expression data (Figure 2C), the number of *mpeg1*+ macrophages in *csf1ra*^V614M^ CHT did not differ from those in WT (Supplemental Figure S2D), however, it was significantly decreased in *csf1rb*^Δ4bp^ fish (Supplemental Figure S2E). To reveal the Csf1a ligand-receptor specificity, we microinjected *csf1a* into both of these mutants and we demonstrate that ligand-overexpression induced macrophage expansion was defective in them. Neither the number of *mpeg1:EGFP*+ cells in *csf1ra*^V614M^ (Supplemental Figure S2F) nor of *mpeg1+* cells in *csf1rb*^Δ4bp^ mutants (Supplemental Figure S2G) was changed as compared to the WT. Thus, Csf1a acts through both Csf1rs.

We examined *lcp1* expression in *csf1a* and *csf1b* ligand mutants, carrying frameshift mutations. ISH showed a significant decrease in the number of *lcp1* expressing cells in the *csf1a^ins2bp^* but not in the *csf1b^Δ2bp^* mutants (Figure 2C). Likewise, this phenotype can be rescued at 36 hpf by injection of *csf1a* mRNA or Csf1a proteins (Supplemental Figure S3A, B) into *csf1a*^ins2bp^ mutant 1-cell stage embryos (Figure 2D).

### Zebrafish *csf1rb* together with *il34* regulate embryonic granulocytic fates

To test whether *csf1r* signaling is involved in the generation of other myeloid cell types besides macrophages, we examined the granulocytic lineage in *csf1* ligand and receptor mutants. Mature granulocytes were visualized in zebrafish embryos and larvae by Sudan black B (SBB) staining and positive cells were counted in tails (Figure 3A).

**Figure 3:**
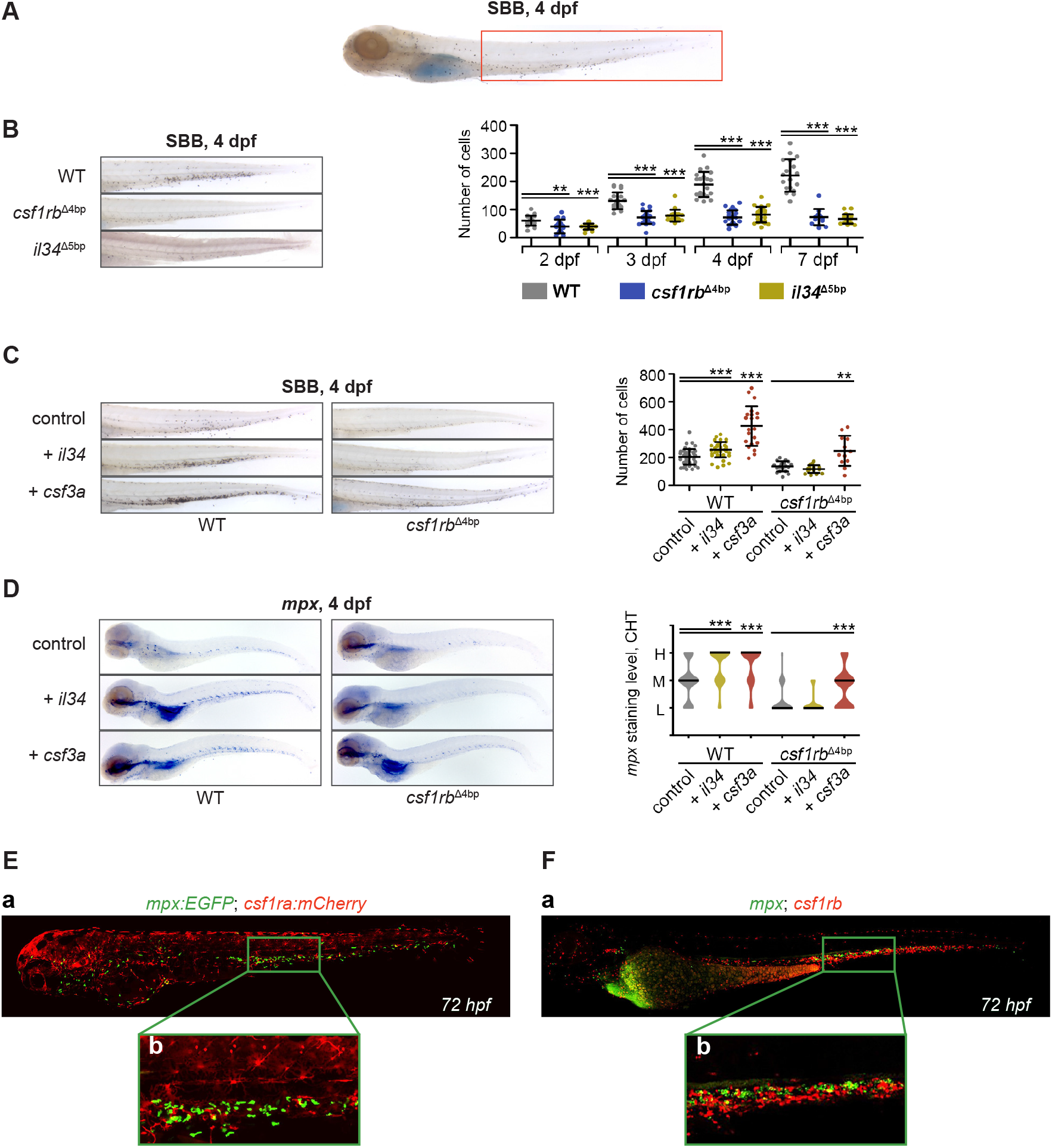
*il34* binds to *csf1rb* and regulates the embryonic granulocytic fate: (A) Sudan black B (SBB) staining of a 4 dpf larva. The analyzed area is marked with a red rectangle. (B-C) SBB positive cells were manually counted and the level of statistical significance was determined by unpaired two-tailed t-test. **P < 0.006, ***P < 0.0001. (B) SBB in WT, *csf1rb^Δ4bp^* and *il34^Δ5bp^* at 4 dpf. The graph on the right shows the number of SBB positive cells during zebrafish embryonal and larval development, 2 – 7 dpf. (C) *il34* and *csf3a* ligands were overexpressed by mRNA microinjection in 1-cell stage WT or *csf1rb^Δ4bp^* mutant embryos. Control embryos were injected with PBS. SBB staining was performed at 4 dpf. The graph on the right shows the number of SBB positive cells. (D) WISH of 4 dpf larvae showing the expression of *mpx* in WT or mutant *csf1rb^Δ4bp^* embryos with overexpressed *il34* or *csf3a* ligands. Violin plots show the level of *mpx* expression in individual embryos (L = low, M = medium, H = high) with median represented by a black line. ***P < 0.001. (E) Co-expression of *csf1ra* (red) and *mpx* (green) visualized in 72 hpf double transgenic embryos Tg(*fms:GAL4;UAS:mCherry*);Tg(*mpx:EGFP*): (Ea) whole embryo, (Eb) caudal hematopoietic tissue (CHT) region. (F) HCR WISH of 72 hpf embryos for *csf1rb* (red) and *mpx* (green). (Fa) whole embryo, (Fb) CHT region. All SBB staining and WISH bright field images were acquired on Zeiss Axio Zoom.V16 with Zeiss Axiocam 105 color camera and processed using the Extended Depth of Focus module in the ZEN Blue software. FIJI and Adobe Photoshop were used for image processing. Fluorescence images were taken on Dragonfly 503 microscope (Andor) using Zyla 4.2 PLUS sCMOS camera, with 10x magnification and processed with the Fusion software, FIJI and Adobe Photoshop.

Strikingly, the number of SBB+ granulocytes in the tail was lower comparing to the WT and failed to increase during the development in *csf1rb*^Δ4bp^ as well as in *il34*^Δ5bp^ mutants (Figure 3B), whereas it gradually increased in both *csf1ra* mutants at the same time (Supplemental Figure S4A). An intermediate phenotype was documented in the receptor double mutants, in which the size of the original granulocytic pool did not change significantly with time (Supplemental Figure S4A). In addition, the mutation in csf1a or csf1b ligands had no obvious effect on the number of granulocytes (data not shown). Along with these findings, the expression of *mpx* was also significantly downregulated in the CHT region of *csf1rb*^Δ4bp^ and *il34^Δ5bp^*, but not in other receptor or ligand mutants at 4 dpf (Supplemental figure S4B and data not shown).

Further, we assessed the effects of *il34* injection on granulocytic expansion in WT as well as in *csf1rb*^Δ4bp^ mutants. As a positive control, we injected *colony stimulating factor 3a* (*csf3a,* also known as *gcsfa*).^60^ As expected, the injection of *csf3a* mRNA led to a significant increase of SBB positive granulocytes in either *csf1rb*^Δ4bp^ mutant or WT fish at 4 dpf. Similarly, the injection of *il34* mRNA into WT fish also caused an increase, however in contrast to *csf3a*, this *il34* mediated phenotype was diminished in the *csf1rb*^Δ4bp^ mutants (Figure 3C). The same effect was confirmed by *mpx* ISH of 4 dpf injected WT as well as *csf1rb*^Δ4bp^ mutant embryos (Figure 3D). We also tested the other ligands, *csf1a* and *csf1b*, but *il34* was the only one to affect granulocytic expansion (Supplemental Figure S4C). Importantly, microinjection of *il34* induced granulocytic expansion in the *csf1ra*^V614M^ mutant (Supplemental Figure S4D). The co-expression of *mpx* with *csf1rs* in the CHT of 72 hpf embryos shows basically no overlap between *csf1ra* and *mpx* (Figure 3Ea, 3Eb). However, there is a small proportion of *csf1rb* and *mpx* double positive cells (Figure 3Fa, 3Fb). Taken together, these results indicate that Il34 regulates embryonic granulocyte development through Csf1rb.

### Zebrafish *csf1rb* is indispensable for definitive granulopoiesis

To investigate the importance of *csf1rb* in adult granulopoiesis, we imaged and counted the number of *mpx*+ cells in tail fins of 6 months old *mpx:EGFP* transgenic WT and *csf1rb*^Δ4bp^ mutant animals. There was a significantly reduced number of granulocytes in the periphery of *csf1rb* ^Δ4bp^ mutants (Figure 4A). Furthermore, we examined WKMs of *csf1ra*^V614M^*, csf1ra*^Δ5bp^*, csf1rb*^Δ4bp^ *and il34*^Δ5bp^ animals using FACS analysis (Figure 4B), noticing a significant decrease in the number of myeloid cells in *csf1rb*^Δ4bp^ mutants only (Mean ± SD; WT: 39.7 ± 3.2%; *csf1rb*^Δ4bp^: 16.9 ± 6.4%; *il34*^Δ5bp^: 48.3 ± 1.7%). The other mutants were not affected (data not shown). Additionally, we prepared thin layer smears stained by May-Grünwald and Giemsa (MGG) from WKM cell suspensions (Supplemental Figure S5A). Most of the cells isolated from *csf1rb*^Δ4bp^ mutants resembled immature undifferentiated cells. The morphology of *csf1rb*^Δ4bp^ mutant granulocytes was abnormal with a significantly decreased frequency of lobulated mature cells that were much smaller in size compared to WT cells.

**Figure 4:**
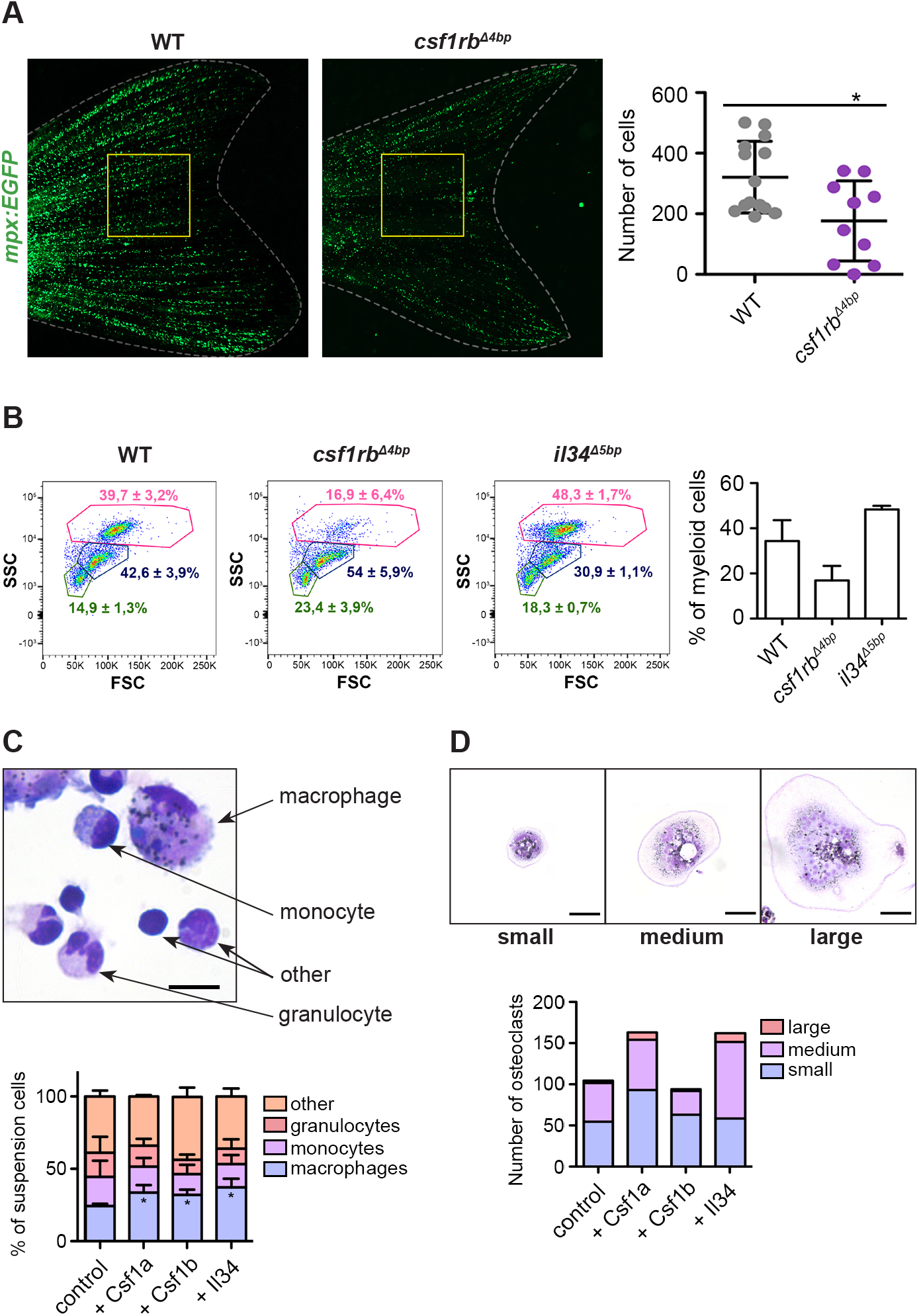
*csf1rb* is indispensable for definitive granulopoiesis. (A) Number of neutrophils in adult Tg(*mpx:EGFP*) = WT and Tg(*mpx:EGFP*);*csf1rb^Δ4bp^* = *csf1rb^Δ4bp^* fish tails. Neutrophils were manually counted in the area of the yellow square. The level of statistical significance was determined by unpaired two-tailed t-test. *P < 0,04. (B) FACS analysis of whole kidney marrow (WKM) cell suspension from WT, *csf1rb^Δ4bp^* and *il34^Δ5bp^* adult zebrafish. The numbers in FSC/SSC plots represent the mean percentage with SD in the gates of myeloid cells (pink gate), progenitors (blue gate) and lymphoid and small progenitor cells (green gate). The percentage of WKM cells in the myeloid gate is also shown in the bar graph on the right. (C-D) *Ex vivo* culture of WKM cells treated with Csf1a, Csf1b or Il34 proteins. (C) After 3 days in culture, smears of suspension cells were stained on microscopic glass slides with May-Grünwald and Giemsa (MGG) and the number of differentiated cells (monocytes, macrophages and granulocytes) was counted. The graph on the bottom shows the mean percentage of cells with SD. *P < 0,04. The scale bar on the microscopic image is 20 μm. (D) After 3 days in culture, adherent cells on the dish were washed with PBS, stained with MGG and the number of small, medium and large osteoclasts was counted in 20 fields of view with a 20x magnification objective. The scale bar on the microscopic image is 50 μm. Fluorescence images were acquired on Zeiss Axio Zoom.V16 with Zeiss Axiocam 506 mono camera and ZEN Blue software. ImageJ and Adobe Photoshop were used for image processing. Bright field images of *ex vivo* cultures were acquired on Leica DM 2000 microscope with Zeiss Axiocam 105 color camera.

### Csf1a, Csf1b and Il34 zebrafish proteins expand adult myeloid cells in *ex vivo* culture

To investigate cell autonomous effects of Csf1 and Il34 cytokines, we performed *in vitro* experiments using recombinant ligand proteins. Therefore, we isolated and seeded WKM cells from WT fish, as published previously,^56,60^ with the addition of recombinant zebrafish Csf1a, Csf1b or Il34 proteins. After 3 days in suspension culture, we prepared histological smears of myeloid cells for enumeration. Specifically, we counted the number of monocytes, differentiated macrophages and granulocytes in proportion to other cells (mostly immature blood progenitors and lymphoid-like cells). In the presence of any of all three cytokines, suspension cells differentiated towards the myeloid lineage to mostly become mature macrophages (Figure 4C). Strikingly, mature multinucleated osteoclasts represented a major fraction of adherent cells. The addition of Csf1a or Il34 to the *ex vivo* culture, promoted the proliferation of osteoclast progenitors and their fusion. (Figure 4D).

### Myelopoiesis is partially blocked in the *csf1rb*^Δ4bp^ mutants

Our results thus far have shown that embryonic granulopoiesis is altered in *csf1rb*^Δ4bp^ mutants. With the noted differences in composition of individual hematopoietic population between WT and mutant animals, we were interested in characterizing these changes at the single cell level. We thus utilized scRNA-seq to profile WKM cells isolated from 12 months old WT, *csf1ra*^Δ5bp^ and *csf1rb*^Δ4bp^ mutants.

Via unsupervised clustering of single cell transcriptomes and based upon known lineage marker genes, we named each cluster based on likely cell type origins (Figure 5A). The percentage of cells in selected clusters of blood progenitors (BP), myeloid progenitors (MP), monocytes/macrophages (M/M) and granulocytes (G) is shown in a table (Figure 5B). We saw increased numbers of cells in BP and MP populations for *csf1ra*^Δ5bp^ and surprisingly G for *csf1rb*^Δ4bp^ (Figure 5B, Supplemental Figure S6A), whereas the number of M/M was decreased in *csf1rb*^Δ4bp^ as expected.

**Figure 5:**
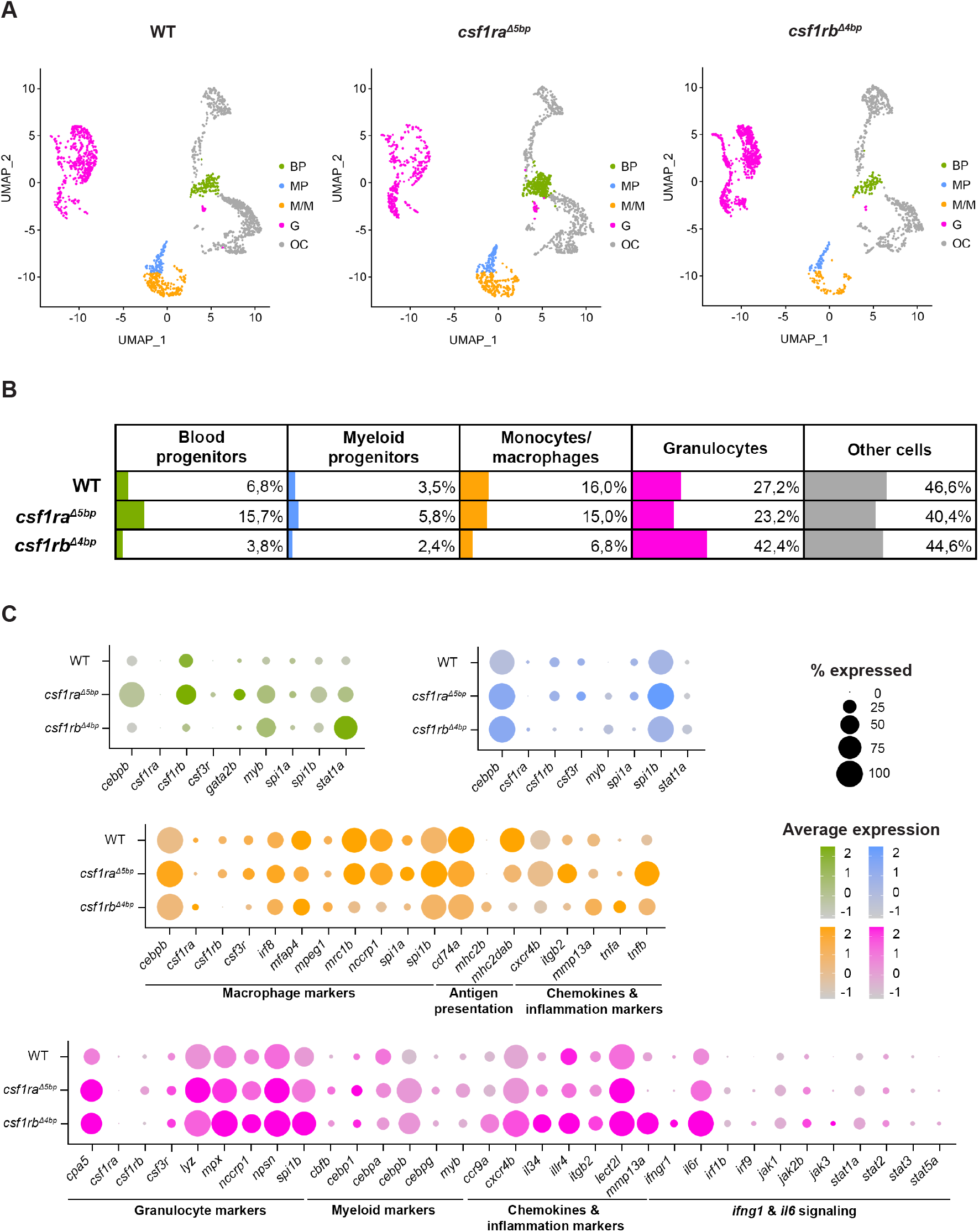
Single-cell RNA sequencing (scRNA-seq) of adult WT, *csf1ra*^Δ5bp^ and *csf1rb*^Δ4bp^ WKM cells shows differentially expressed hematopoietic genes. (A) Clusters in Dim plots represent individual selected populations of WKM hematopoietic cells. The identity of each cluster is based on likely cell origin according to lineage marker gene expression. Green – blood progenitors (BP), blue – myeloid progenitors (MP), orange – monocytes/macrophages (M/M), pink – granulocytes (G), gray – other cells (OC). (B) Table representing the percentage of cells in each population for WT and respective mutants. (C) Dot plot visualization of scRNA-seq gene expression in individual populations of WKM hematopoietic cells of WT and respective mutants. The color of each dot represents the level of expression (also depicted in the histogram) and the size of the dot is showing the percentage of cells expressing each individual gene. Green – BP, blue – MP, orange – M/M, pink – G.

Further, we picked out representative markers of hematopoiesis that characterize BP (green), MP (blue), M/M (orange), and G (pink) populations and created dot plot expression schemes showing their relative expression and the percent of cells expressing them (Figure 5C). We observed deregulation in most of these markers in both mutants. However, for *csf1ra*^Δ5bp^ mutant the differences were more prominent in progenitors (BP and MP) and for *csf1rb*^Δ4b^ they were more prominent in differentiated myeloid cells (M/M and G). Additionally, we observed that the *csf1rb* gene was upregulated in *csf1ra*^Δ5bp^ mutant and *vice versa*. After a closer look at the discrepancy to the decreased number of embryonic granulocytes and the peripheral neutropenia observed in adult *csf1rb*^Δ4bp^ fish, we noted that a high proportion of granulocyte-like cells accumulate in WKM (Supplemental Figure S6A) and there are aberrantly expressed progenitor (Supplemental Figure S6B), migration as well as inflammation markers (Figure 5C, Supplemental Figure S6C).

## Discussion

Differentiation, survival and maturation of myeloid cells is tightly controlled by extrinsic factors, such as cytokines. To the most prominent ones belong CSF1, CSF2 (also known as granulocyte-macrophage-CSF, GM-CSF), and CSF3. The critical role of CSF1 signaling on proper macrophage cell differentiation and survival was shown in mice mutants that lack *CSF1R* or *CSF1*.^9,61,62^ Interestingly, *CSF1*, *CSF2* and *CSF3* triple mutant mice are not completely devoid of macrophages or granulocytes.^62^ This indicates that the role of these factors can be replaced by other cytokines, such as IL-6 or IL-34.^4,63,64^ Although CSF signaling has been historically studied mainly in mammalian and bird animal models and many transgenic and mutant lines are available, zebrafish has recently become a popular alternative model organism for modeling hematopoietic human diseases.

Here, we studied the *in vivo* function of zebrafish Csf1 receptor paralogs (*csf1ra*, *csf1rb*) and their ligands (Csf1a, Csf1b, Il34) to describe their involvement in fish myelopoiesis and to characterize their diversification. Our findings demonstrate that Csf1a and Csf1b sub-functionalized during embryonic myelopoiesis. Specifically, we show that only the Csf1a ligand is important in the development of embryonic macrophages, while both, Csf1a and Csf1b, are involved in pigmentation. This is consistent with previous studies demonstrating the role of Csf1a/b in adult pigment patterning.^15,65^

To get detailed insight into the involvement of Csf1 signaling in embryonic and adult macrophage development in zebrafish, we decided to study Csf1 ligand and receptor mutants. Zebrafish *csf1ra, csf1rb* and *il34* mutants have been previously described for the lack of microglia.^42,49^ It was shown that macrophages develop, migrate or proliferate aberrantly in *il34*^Δ5bp^ mutants,^43,49^ and in *csf1ra*^V614M^;*csf1rb*^Δ4bp^ double mutant fish,^66^ whereas they develop normally in *csf1ra*^V614M^ (known as Panther)^42,43^ and *csf1rb*^sa1503^ mutant^44^ embryos. The status of macrophage development in *csf1rb*^Δ4bp^ single mutant animals has remained unnoticed.

In contrast to these studies, we demonstrate that the number of embryonic macrophages in the CHT of *csf1ra*^Δ5bp^ and *csf1rb*^Δ4bp^ mutant embryos is decreased. We suppose that the discrepancy in the number of embryonic macrophages between *csf1ra*^V614M^ and *csf1ra*^Δ5bp^ or *csf1rb*^sa1503^ and *csf1rb*^Δ4bp^ could be explained by the fact that *csf1ra*^Δ5bp^ and *csf1rb*^Δ4bp^ mutants have stronger phenotypes because they carry frameshift mutations with a premature stop codon. This has been also shown in another Panther mutant with a C-terminal deletion mutation (*csf1ra*^j4blue^).^48,67,68^ Surprisingly, macrophage expansion is diminished in both *csf1ra*^V614M^ and *csf1rb*^Δ4bp^ mutants after *csf1a* microinjection. Therefore, we hypothesize that Csf1a drives macrophage development via either of both, Csf1ra or Csf1rb, receptors.

Besides the role of Csf1 signaling in macrophage differentiation, we noticed that it might also be involved in zebrafish granulocyte differentiation. Despite decades of CSF1R research, only sparse evidence exists linking CSF1R signaling to granulocyte differentiation in mammals.^69^ Strikingly, we show that *csf1rb*^Δ4bp^ and also *il34*^Δ5bp^ mutant zebrafish embryos have major defects in granulopoiesis and that the overexpression of *il34* in WT animals leads to granulocyte expansion. Further, we show that the pool of embryonic granulocytes cannot be expanded in the *csf1rb*^Δ4bp^ mutants by *il34*. This implies that Il34 plays a previously unknown role in the regulation of embryonic granulopoiesis and that it can act through Csf1rb, providing a novel Csf3r alternative pathway that is important for granulocyte differentiation.

As a next step, we decided to examine adult macrophages in receptor mutant animals. In contrast to defects in embryonic macrophage myelopoiesis, we did not see any evident myeloid defects in WKM of adult *csf1ra*^Δ5bp^ mutants when examined histologically and by FACS. Nevertheless, the scRNA-seq analysis of *csf1ra*^Δ5bp^ mutants revealed that there is a slight downregulation of macrophage-specific markers, such as *mfap4*, *mhc2dab*, or *mrc1b* and an upregulation of progenitor specific markers, such as *myb*, *cebpa* and *gata2b* and the overall number of blood and myeloid progenitors is elevated as well, however, the other populations are mostly unchanged. Despite these less severe phenotypes in adult WKM cells, we hypothesize that besides Csf1ra role in development of embryonic macrophages, it also plays a minor role during adult hematopoiesis as previously not shown. The decreased number of monocytes/macrophages was also observed in *csf1rb*^Δ4bp^ mutant using scRNA-seq. In addition, markers specific to macrophage host defense/phagosome and antigen presentation are decreased (*mrc1b, nccrp1, mhc2dab*) in this mutant. These observations led us to the conclusion that Csf1rb is equally important during embryonic as well as adult macrophage development.

Additionally, *csf1rb*^Δ4bp^ adult fish have lower number of myeloid cells in WKM, and also fewer *mpx*+ cells in the periphery. Since the majority of myeloid cells in WKM are neutrophils^70,71^ we attribute the reduced number of myeloid cells to the loss of mature, physiologically normal granulocytes. However, surprisingly our scRNA-seq data indicate that *csf1rb*^Δ4bp^ mutants have more granulocytes in WKM. We explain this discrepancy by the fact that *csf1rb*^Δ4bp^ granulocyte progenitors cannot fully differentiate and migrate, and progenitor-like early and also late granulocytes accumulate in WKM. This is proven by scRNA-seq marker gene expression profiling and also by FACS and histological staining of WKM cells, demonstrating that *csf1rb*^Δ4bp^ fish have a high proportion of small cells with abnormal morphology and low granularity. Using scRNA-seq analysis, the predicted granulocyte population of *csf1rb*^Δ4bp^ WKMs showed an overexpression of *myb, cbfb, spi1b* and *cebp* markers, known to play essential roles in progenitor hemostasis.^72–76^ Further we observed deregulated expression of chemokine and inflammation markers (*il34*, *itgb2*, *mmp13a*, *il6r*, *ifngr1*), and markers connected to granulocyte migration (*cxcr4b*, *ifngr1, cxcl8b*).^77,78^ Abnormal activation of the zebrafish *ifngr1* signaling has been previously shown in neutropenic zebrafish.^79^

Next, we have shown that *csf1r* paralogues have functionally diverged in the course of teleost evolution. Consistently with previous findings,^44^ we suggest that the function and expression of *csf1* receptor paralogues is mostly non-overlapping. At the single cell level, using scRNA-seq analysis of adult WKM cells, we showed that there is no expression overlap between *csf1ra* and *csf1rb* and that *csf1rb* is highly expressed in blood and myeloid progenitors as well as in a small subset of monocytes/macrophages and granulocytes. In contrast, the *csf1ra* expression is restricted only to a small subset of monocytes/macrophages. In correlation with these expression data, we also demonstrated that mutation in *csf1ra* primarily affects embryonic macrophage development, while the mutation in *csf1rb* is equally important during embryonic as well as adult development of both macrophages and granulocytes. Regarding ligand-receptor signaling specificities, we found that Csf1a acts via both, Csf1ra and Csf1rb and that the granulocyte related function of Il34 is executed via Csf1rb. In contrast, Csf1b has no function during embryonic myelopoiesis and it expands xanthophores instead, together with Csf1a. Importantly, based on the fact that there is low expression of *csf1rb* in the skin, we assume that the Csf1 dependent xanthophore expansion is mediated via Csf1ra.

In addition, our *ex vivo* experiments suggest that recombinant Csf1a/b and Il34 cytokines are functional in *ex vivo* cultures and drive WKM-derived myeloid cell differentiation towards monocyte-macrophage and osteoclast fates. However, terminal granulocytic differentiation is not affected by Csf1r ligands.

In summary, we demonstrate that zebrafish continues to provide new biological insights relevant to disease. Using a wide range of convenient *in vivo* and *ex vivo* tools, it is possible to characterize new exciting roles of cytokines under steady-state as well as non-steady-state conditions. Myeloid cells, including neutrophils and macrophages are critical actors in cancerogenesis^80–82^ and the CSF1 pathway is a promising target for clinical treatments.^24,83^ Here we performed detailed characterization of Csf1 signaling in zebrafish making it suitable for preclinical disease modeling in high-throughput discovery of new therapeutics.

## Acknowledgements

We thank Nikol Pavlu, Tereza Hojerova and Tereza Hingarova for animal care, Trevor Epp for editing the manuscript, and Leonard Zon for providing *mpeg1:EGFP* and *mpx:EGFP* reporter fish lines. We acknowledge Michal Kolar for help with sc-RNA-transcriptomics (LM2018131). We acknowledge the Light Microscopy Core Facility, IMG CAS, Prague, Czech Republic, supported by MEYS (LM2018129, CZ.02.1.01/0.0/0.0/18_046/0016045) for their support with the confocal imaging. This work was supported by the Czech Science Foundation (18-18363S), LM2018131 and 68378050-KAV-NPUI to PB and by the ERC Advanced Grant “DanioPattern” (694289) to C.N.-V. and U.I.

## Authorship

Contribution: M.H., T.M., O.M., A.P., U.I. performed research. O.S. and P.B. designed the research. T.J.v.H. and C.N.-V. provided critical reagents for the work. M.H., T.M., P.B. and O.S. wrote the manuscript.

## References

1. Knapp DJ, Hammond CA, Aghaeepour N, et al. Distinct signaling programs control human hematopoietic stem cell survival and proliferation. Blood. 2017;129(3):307–318.

2. Buenrostro JD, Corces MR, Lareau CA, et al. Integrated Single-Cell Analysis Maps the Continuous Regulatory Landscape of Human Hematopoietic Differentiation. Cell. 2018;173(6):1535–1548 e1516.

3. Chen X, Liu H, Focia PJ, Shim AH, He X. Structure of macrophage colony stimulating factor bound to FMS: diverse signaling assemblies of class III receptor tyrosine kinases. Proc Natl Acad Sci U S A. 2008;105(47):18267–18272.

4. Chihara T, Suzu S, Hassan R, et al. IL-34 and M-CSF share the receptor Fms but are not identical in biological activity and signal activation. Cell Death Differ. 2010;17(12):1917–1927.

5. Lin W, Xu D, Austin CD, et al. Function of CSF1 and IL34 in Macrophage Homeostasis, Inflammation, and Cancer. Front Immunol. 2019;10:2019.

6. Bencheikh L, Diop MK, Riviere J, et al. Dynamic gene regulation by nuclear colony-stimulating factor 1 receptor in human monocytes and macrophages. Nat Commun. 2019;10(1):1935.

7. Liu H, Leo C, Chen X, et al. The mechanism of shared but distinct CSF-1R signaling by the non-homologous cytokines IL-34 and CSF-1. Biochim Biophys Acta. 2012;1824(7):938–945.

8. Wiktor-Jedrzejczak W, Bartocci A, Ferrante AW, Jr., et al. Total absence of colony-stimulating factor 1 in the macrophage-deficient osteopetrotic (op/op) mouse. Proc Natl Acad Sci U S A. 1990;87(12):4828–4832.

9. Dai XM, Ryan GR, Hapel AJ, et al. Targeted disruption of the mouse colony-stimulating factor 1 receptor gene results in osteopetrosis, mononuclear phagocyte deficiency, increased primitive progenitor cell frequencies, and reproductive defects. Blood. 2002;99(1):111–120.

10. Van Wesenbeeck L, Odgren PR, MacKay CA, et al. The osteopetrotic mutation toothless (tl) is a loss-of-function frameshift mutation in the rat Csf1 gene: Evidence of a crucial role for CSF-1 in osteoclastogenesis and endochondral ossification. Proc Natl Acad Sci U S A. 2002;99(22):14303–14308.

11. Garcia-Morales C, Rothwell L, Moffat L, et al. Production and characterisation of a monoclonal antibody that recognises the chicken CSF1 receptor and confirms that expression is restricted to macrophage-lineage cells. Dev Comp Immunol. 2014;42(2):278–285.

12. Hume DA, Gutowska-Ding MW, Garcia-Morales C, et al. Functional evolution of the colony-stimulating factor 1 receptor (CSF1R) and its ligands in birds. J Leukoc Biol. 2020;107(2):237–250.

13. Wang T, Kono T, Monte MM, et al. Identification of IL-34 in teleost fish: differential expression of rainbow trout IL-34, MCSF1 and MCSF2, ligands of the MCSF receptor. Mol Immunol. 2013;53(4):398–409.

14. Hume DA, Irvine KM, Pridans C. The Mononuclear Phagocyte System: The Relationship between Monocytes and Macrophages. Trends Immunol. 2019;40(2):98–112.

15. Caetano-Lopes J, Henke K, Urso K, et al. Correction: Unique and non-redundant function of csf1r paralogues in regulation and evolution of post-embryonic development of the zebrafish. Development. 2020;147(10).

16. Liu W, Di Q, Li K, et al. The synergistic role of Pu.1 and Fms in zebrafish osteoclast-reducing osteopetrosis and possible therapeutic strategies. J Genet Genomics. 2020.

17. Rademakers R, Baker M, Nicholson AM, et al. Mutations in the colony stimulating factor 1 receptor (CSF1R) gene cause hereditary diffuse leukoencephalopathy with spheroids. Nat Genet. 2011;44(2):200–205.

18. Oosterhof N, Holtman IR, Kuil LE, et al. Identification of a conserved and acute neurodegeneration-specific microglial transcriptome in the zebrafish. Glia. 2017;65(1):138–149.

19. Oosterhof N, Chang IJ, Karimiani EG, et al. Homozygous Mutations in CSF1R Cause a Pediatric-Onset Leukoencephalopathy and Can Result in Congenital Absence of Microglia. Am J Hum Genet. 2019;104(5):936–947.

20. Hagan N, Kane JL, Grover D, et al. CSF1R signaling is a regulator of pathogenesis in progressive MS. Cell Death Dis. 2020;11(10):904.

21. Boultwood J, Rack K, Kelly S, et al. Loss of both CSF1R (FMS) alleles in patients with myelodysplasia and a chromosome 5 deletion. Proc Natl Acad Sci U S A. 1991;88(14):6176–6180.

22. Gu TL, Mercher T, Tyner JW, et al. A novel fusion of RBM6 to CSF1R in acute megakaryoblastic leukemia. Blood. 2007;110(1):323–333.

23. Roh-Johnson M, Shah AN, Stonick JA, et al. Macrophage-Dependent Cytoplasmic Transfer during Melanoma Invasion In Vivo. Dev Cell. 2017;43(5):549–562 e546.

24. Durham BH, Lopez Rodrigo E, Picarsic J, et al. Activating mutations in CSF1R and additional receptor tyrosine kinases in histiocytic neoplasms. Nat Med. 2019;25(12):1839–1842.

25. Hume DA, Caruso M, Ferrari-Cestari M, Summers KM, Pridans C, Irvine KM. Phenotypic impacts of CSF1R deficiencies in humans and model organisms. J Leukoc Biol. 2019.

26. Guillonneau C, Bézie S, Anegon I. Immunoregulatory properties of the cytokine IL-34. Cellular and Molecular Life Sciences. 2017;74(14):2569–2586.

27. Mantovani A, Marchesi F, Malesci A, Laghi L, Allavena P. Tumour-associated macrophages as treatment targets in oncology. Nat Rev Clin Oncol. 2017;14(7):399–416.

28. Peranzoni E, Donnadieu E. Improving efficacy of cancer immunotherapy through targeting of macrophages. Hum Vaccin Immunother. 2019;15(1):189–192.

29. Cannarile MA, Weisser M, Jacob W, Jegg AM, Ries CH, Ruttinger D. Colony-stimulating factor 1 receptor (CSF1R) inhibitors in cancer therapy. J Immunother Cancer. 2017;5(1):53.

30. Czako B, Marszalek JR, Burke JP, et al. Discovery of IACS-9439, a Potent, Exquisitely Selective, and Orally Bioavailable Inhibitor of CSF1R. J Med Chem. 2020;63(17):9888–9911.

31. Ransom DG, Haffter P, Odenthal J, et al. Characterization of zebrafish mutants with defects in embryonic hematopoiesis. Development. 1996;123:311–319.

32. Howe K, Clark MD, Torroja CF, et al. The zebrafish reference genome sequence and its relationship to the human genome. Nature. 2013;496(7446):498–503.

33. Britto DD, Wyroba B, Chen W, et al. Macrophages enhance Vegfa-driven angiogenesis in an embryonic zebrafish tumour xenograft model. Dis Model Mech. 2018;11(12).

34. Kimmel CB, Ballard WW, Kimmel SR, Ullmann B, Schilling TF. Stages of embryonic development of the zebrafish. Dev Dyn. 1995;203(3):253–310.

35. Yan C, Brunson DC, Tang Q, et al. Visualizing Engrafted Human Cancer and Therapy Responses in Immunodeficient Zebrafish. Cell. 2019.

36. Fazio M, Ablain J, Chuan Y, Langenau DM, Zon LI. Zebrafish patient avatars in cancer biology and precision cancer therapy. Nat Rev Cancer. 2020;20(5):263–273.

37. Xiao J, Glasgow E, Agarwal S. Zebrafish Xenografts for Drug Discovery and Personalized Medicine. Trends Cancer. 2020;6(7):569–579.

38. Ablain J, Liu S, Moriceau G, Lo RS, Zon LI. SPRED1 deletion confers resistance to MAPK inhibition in melanoma. J Exp Med. 2021;218(3).

39. Taylor JS, Braasch I, Frickey T, Meyer A, Van de Peer Y. Genome duplication, a trait shared by 22000 species of ray-finned fish. Genome Res. 2003;13(3):382–390.

40. Oltova J, Svoboda O, Bartunek P. Hematopoietic Cytokine Gene Duplication in Zebrafish Erythroid and Myeloid Lineages. Front Cell Dev Biol. 2018;6:174.

41. Pazhakh V, Lieschke GJ. Hematopoietic growth factors: the scenario in zebrafish. Growth Factors. 2018;36(5-6):196–212.

42. Oosterhof N, Kuil LE, van der Linde HC, et al. Colony-Stimulating Factor 1 Receptor (CSF1R) Regulates Microglia Density and Distribution, but Not Microglia Differentiation In Vivo. Cell Rep. 2018;24(5):1203–1217 e1206.

43. Wu S, Xue R, Hassan S, et al. Il34-Csf1r Pathway Regulates the Migration and Colonization of Microglial Precursors. Dev Cell. 2018;46(5):552–563 e554.

44. Ferrero G, Miserocchi M, Di Ruggiero E, Wittamer V. A csf1rb mutation uncouples two waves of microglia development in zebrafish. Development. 2020.

45. Ginhoux F, Jung S. Monocytes and macrophages: developmental pathways and tissue homeostasis. Nat Rev Immunol. 2014;14(6):392–404.

46. Alestrom P, D’Angelo L, Midtlyng PJ, et al. Zebrafish: Housing and husbandry recommendations. Lab Anim. 2020;54(3):213–224.

47. Oltova J, Jindrich J, Skuta C, Svoboda O, Machonova O, Bartunek P. Zebrabase: An Intuitive Tracking Solution for Aquatic Model Organisms. Zebrafish. 2018;15(6):642–647.

48. Parichy DM, Ransom DG, Paw B, Zon LI, Johnson SL. An orthologue of the kit-related gene fms is required for development of neural crest-derived xanthophores and a subpopulation of adult melanocytes in the zebrafish, Danio rerio. Development. 2000;127(14):3031–3044.

49. Kuil LE, Oosterhof N, Geurts SN, van der Linde HC, Meijering E, van Ham TJ. Reverse genetic screen reveals that Il34 facilitates yolk sac macrophage distribution and seeding of the brain. Dis Model Mech. 2019;12(3).

50. Ellett F, Pase L, Hayman JW, Andrianopoulos A, Lieschke GJ. mpeg1 promoter transgenes direct macrophage-lineage expression in zebrafish. Blood. 2011;117(4):E49–E56.

51. Gray C, Loynes CA, Whyte MK, Crossman DC, Renshaw SA, Chico TJ. Simultaneous intravital imaging of macrophage and neutrophil behaviour during inflammation using a novel transgenic zebrafish. Thromb Haemost. 2011;105(5):811–819.

52. Renshaw SA, Loynes CA, Trushell DM, Elworthy S, Ingham PW, Whyte MK. A transgenic zebrafish model of neutrophilic inflammation. Blood. 2006;108(13):3976–3978.

53. Seger C, Hargrave M, Wang X, Chai RJ, Elworthy S, Ingham PW. Analysis of Pax7 expressing myogenic cells in zebrafish muscle development, injury, and models of disease. Dev Dyn. 2011;240(11):2440–2451.

54. Choi HM, Calvert CR, Husain N, et al. Mapping a multiplexed zoo of mRNA expression. Development. 2016;143(19):3632–3637.

55. Schindelin J, Arganda-Carreras I, Frise E, et al. Fiji: an open-source platform for biological-image analysis. Nature Methods. 2012;9(7):676–682.

56. Svoboda O, Stachura DL, Machonova O, Zon LI, Traver D, Bartunek P. Ex vivo tools for the clonal analysis of zebrafish hematopoiesis. Nat Protoc. 2016;11(5):1007–1020.

57. Fleming SJ, Marioni JC, Babadi M. CellBender remove-background: a deep generative model for unsupervised removal of background noise from scRNA-seq datasets. bioRxiv. 2019;791699.

58. Stuart T, Butler A, Hoffman P, et al. Comprehensive Integration of Single-Cell Data. Cell. 2019;177(7):1888–1902 e1821.

59. Mahalwar P, Walderich B, Singh AP, Nusslein-Volhard C. Local reorganization of xanthophores fine-tunes and colors the striped pattern of zebrafish. Science. 2014;345(6202):1362–1364.

60. Stachura DL, Svoboda O, Campbell CA, et al. The zebrafish granulocyte colony-stimulating factors (Gcsfs): 2 paralogous cytokines and their roles in hematopoietic development and maintenance. Blood. 2013;122(24):3918–3928.

61. Dai X-M, Zong X-H, Sylvestre V, Stanley ER. Incomplete restoration of colony-stimulating factor 1 (CSF-1) function in CSF-1–deficient Csf1op/Csf1op mice by transgenic expression of cell surface CSF-1. Blood. 2004;103(3):1114–1123.

62. Hibbs ML, Quilici C, Kountouri N, et al. Mice lacking three myeloid colony-stimulating factors (G-CSF, GM-CSF, and M-CSF) still produce macrophages and granulocytes and mount an inflammatory response in a sterile model of peritonitis. J Immunol. 2007;178(10):6435–6443.

63. Lelios I, Cansever D, Utz SG, Mildenberger W, Stifter SA, Greter M. Emerging roles of IL-34 in health and disease. J Exp Med. 2020;217(3).

64. Wang Y, Szretter KJ, Vermi W, et al. IL-34 is a tissue-restricted ligand of CSF1R required for the development of Langerhans cells and microglia. Nat Immunol. 2012;13(8):753–760.

65. Patterson LB, Parichy DM. Interactions with iridophores and the tissue environment required for patterning melanophores and xanthophores during zebrafish adult pigment stripe formation. PLoS Genet. 2013;9(5):e1003561.

66. Kuil LE, Oosterhof N, Ferrero G, et al. Zebrafish macrophage developmental arrest underlies depletion of microglia and reveals Csf1r-independent metaphocytes. Elife. 2020;9.

67. Pagan AJ, Yang CT, Cameron J, et al. Myeloid Growth Factors Promote Resistance to Mycobacterial Infection by Curtailing Granuloma Necrosis through Macrophage Replenishment. Cell Host Microbe. 2015;18(1):15–26.

68. Morales RA, Allende ML. Peripheral Macrophages Promote Tissue Regeneration in Zebrafish by Fine-Tuning the Inflammatory Response. Front Immunol. 2019;10:253.

69. Sasmono RT, Ehrnsperger A, Cronau SL, et al. Mouse neutrophilic granulocytes express mRNA encoding the macrophage colony-stimulating factor receptor (CSF-1R) as well as many other macrophage-specific transcripts and can transdifferentiate into macrophages in vitro in response to CSF-1. J Leukoc Biol. 2007;82(1):111–123.

70. Lieschke GJ, Oates AC, Crowhurst MO, Ward AC, Layton JE. Morphologic and functional characterization of granulocytes and macrophages in embryonic and adult zebrafish. Blood. 2001;98(10):3087–3096.

71. Traver D, Paw BH, Poss KD, Penberthy WT, Lin S, Zon LI. Transplantation and in vivo imaging of multilineage engraftment in zebrafish bloodless mutants. Nat Immunol. 2003;4(12):1238–1246.

72. Dahl R, Walsh JC, Lancki D, et al. Regulation of macrophage and neutrophil cell fates by the PU.1:C/EBPalpha ratio and granulocyte colony-stimulating factor. Nat Immunol. 2003;4(10):1029–1036.

73. Jin H, Li L, Xu J, et al. Runx1 regulates embryonic myeloid fate choice in zebrafish through a negative feedback loop inhibiting Pu.1 expression. Blood. 2012;119(22):5239–5249.

74. Bresciani E, Carrington B, Wincovitch S, et al. CBFbeta and RUNX1 are required at 2 different steps during the development of hematopoietic stem cells in zebrafish. Blood. 2014;124(1):70–78.

75. Jin H, Huang Z, Chi Y, et al. c-Myb acts in parallel and cooperatively with Cebp1 to regulate neutrophil maturation in zebrafish. Blood. 2016;128(3):415–426.

76. Dai Y, Zhu L, Huang Z, et al. Cebpalpha is essential for the embryonic myeloid progenitor and neutrophil maintenance in zebrafish. J Genet Genomics. 2016;43(10):593–600.

77. Summers C, Rankin SM, Condliffe AM, Singh N, Peters AM, Chilvers ER. Neutrophil kinetics in health and disease. Trends Immunol. 2010;31(8):318–324.

78. Robson RL, McLoughlin RM, Witowski J, et al. Differential regulation of chemokine production in human peritoneal mesothelial cells: IFN-gamma controls neutrophil migration across the mesothelium in vitro and in vivo. J Immunol. 2001;167(2):1028–1038.

79. Fan HB, Liu YJ, Wang L, et al. miR-142-3p acts as an essential modulator of neutrophil development in zebrafish. Blood. 2014;124(8):1320–1330.

80. Wang J, Cao Z, Zhang X, et al. Novel Mechanism of Macrophage-Mediated Metastasis Revealed in a Zebrafish Model of Tumor Development. Cancer Res. 2014.

81. Groth C, Hu X, Weber R, et al. Immunosuppression mediated by myeloid-derived suppressor cells (MDSCs) during tumour progression. Br J Cancer. 2019;120(1):16–25.

82. Tulotta C, Stefanescu C, Chen Q, Torraca V, Meijer AH, Snaar-Jagalska BE. CXCR4 signaling regulates metastatic onset by controlling neutrophil motility and response to malignant cells. Sci Rep. 2019;9(1):2399.

83. Edwards DK, V, Watanabe-Smith K, Rofelty A, et al. CSF1R inhibitors exhibit antitumor activity in acute myeloid leukemia by blocking paracrine signals from support cells. Blood. 2019;133(6):588–599.

