## Supplemental Data for "Colony stimulating factor 1 signaling regulates myeloid fates in zebrafish via distinct action of its receptors and ligands"

### Supplemental methods

#### Cloning of constructs and recombinant protein expression

Zebrafish *csf1a*, *csf1b* and *il34* coding sequences as annotated at Ensembl (GRCZ11) were used to design PCR primers (Supplemental Table 1) for amplifying cDNA fragments from adult zebrafish organs. For mRNA production the pCS2+ plasmid was used containing the following fragments 1. *csf1a* - signal peptide (SP) and cytokine-like (CL) domain without the transmembrane (TM) domain; 2. *csf1b* – full length coding sequence; 3. *il34* – full length coding sequence. Recombinant zebrafish Csf1a, Csf1b and Il34 proteins (Supplemental Figure S3) were prepared as described previously.[1,2] cDNA fragments containing only the CL domains without the SP and TM domain for both *csf1a* and *csf1b* were amplified. Full length *il34* sequence was used (Supplemental Figure S3). For recombinant protein expression the PCR amplified fragments were sub-cloned into the pAc-GP67-B vector containing 6xHis and the baculoviral BaculoGold™ (BD Biosciences) expression system was used. Insect sf21 cells were co-transfected by BaculoGold Bright Baculovirus DNA and pAc-His-Csf1a/Csf1b/Il34 (available via Addgene: 168103/168104/168105). The infected cells express the recombinant proteins extracellularly in large amounts. Purification of His-tagged proteins was done on $Ni^{2+}$-NTA agarose columns according to the manufacturer's protocol (Qiagen). The purity of isolated proteins was determined on western blot by a peroxidase-conjugated poly-histidine monoclonal antibody (Sigma) in PBT (1:5000), as described previously.[3]

#### Generation of mutant zebrafish

The CRISPR/Cas system for gene editing was applied as described previously.[4] Briefly, oligonucleotides (Supplemental table 2) were cloned into pDR274 to generate the sgRNA template vectors. sgRNAs were *in vitro* transcribed from these plasmids using the MEGAscript T7 Transcription Kit (Invitrogen). Cas9 protein was expressed in *E. coli* and purified via an N-termial 6xHis- and Twin-StrepTag. The protein (final concentration 350 ng/µl) was mixed with the sgRNA (final concentration 35 ng/µl) in PBS + 300 mM NaCl, 150 mM KCl, and 2-4-nl of the mixture were injected into one-cell stage embryos. The efficiency of indel generation was tested on eight larvae at 1 dpf by PCR (genotyping primers in Supplemental table 2) and sequence analysis. The remaining larvae were raised to adulthood and mature $F_0$ fish carrying indels were outcrossed. In the $F_1$ generation heterozygous carriers of frame-shift mutations were identified and selected to establish homozygous mutant lines.

The newly generated alleles are:

*csf1a*$^{t33ui}$: 2 bp insertion. c.533_534insCC; p.Tyr178fsX25

*csf1b*$^{t34ui}$: 2 bp deletion. c.79_80delCT; p.Ser28fsX25

*csf1ra*$^{t36ui}$: 5 bp deletion.c.1501_1505delGTCGG; p.Val501fsX3

#### Microinjection of mRNA and proteins

Transcription of mRNA was done *in vitro* by the mMessage mMachine SP6 kit (Roche) from linearized pCS2+ vectors containing the sequences of *csf1a* (available via Addgene: 168110), *csf1b* (168111), and *il34* (168112), respectively. 1-cell stage embryos were microinjected with ~5 nl of the injection solution with 100 ng/ul mRNA concentration for each cytokine. This equates to ~500 pg of each mRNA per embryo. Zebrafish recombinant proteins, Csf1a, Csf1b and Il34 were diluted in PBS to 200 ng/ul and a final amount of ~1000 pg was injected into 1-cell stage embryos.

#### Zebrafish fixation, whole-mount *in situ* hybridization (WISH) and image analysis

Zebrafish embryos were staged for 36 hours post fertilization (hpf), 48 hpf, 72 hpf, 96 hpf and 7 days post fertilization (dpf), incubated on ice for 20 minutes and anesthetized by Tricaine (MS-222) prior to fixation. Fixation was done in 4% paraformaldehyde overnight at 4°C.

Digoxigenin-labeled antisense riboprobes for zebrafish *l-plastin* (*lcp1*) and *myeloperoxidase* (*mpx*) were used for WISH and performed according to standard protocols,[5] with slight modifications.[6] Photomicrographs were taken on a Zeiss Axio Zoom.V16 with Zeiss Axiocam 105 color camera and processed using the Extended Depth of Focus module in the ZEN Blue 2.3 software. The embryos were divided into three categories – low, medium, high – according to the number of positive cells in the tail part of the embryo. Results were visualized in GraphPad Prism.

#### Sudan black B staining of embryos and larvae

Sudan black B (SBB) (Sigma-Aldrich) staining was carried out as described before,[7] with slight changes. Specifically, the fixed embryos were incubated in SBB working solution (0.036% SBB in 70% ethanol with 0.1% phenol) for 90 minutes in the dark, extensively washed in 70% ethanol and the pigment was removed by bleaching prior to imaging. The bleaching solution consisted of 0.8% KOH, 0.9% H2O2, and 0.1% Tween 20. Images were taken on Zeiss Axio Zoom.V16 with Zeiss Axiocam 105 color camera and processed using the Extended Depth of Focus module in the ZEN Blue 2.3 software. The number of SBB positive cells was manually counted in the tail part of the embryo using the Cell counter plugin in Fiji.

#### RNA isolation and qRT-PCR

Whole embryos or adult organs were collected into TRI reagent (Sigma, TR 118) and RNA was isolated according to the manufacturer's instructions. Isolated RNA was treated with RQ1 DNase (Promega). cDNA was transcribed using M-MLV reverse transcriptase (Promega). qPCR experiments were performed in triplicate using LightCycler 480 and SYBR Green (Roche). *ef1a* or *mob4* housekeeping genes were used to normalize gene expression. The list of used primers is in Supplemental Table 3. Relative expression of genes was calculated by the $2^{-(Ct[\text{gene of interest}] - Ct[\text{housekeeping gene}])}$ formula. Statistical analysis was determined from 2-7 biological replicates using unpaired two-tailed t-test.

#### FACS analysis

WKM from adult fish was isolated as described previously.[2] Flow cytometry of single-cell suspension in 1x HBSS was performed with a BD FACSymphony. Data analysis was done using BD FACSDiva and FlowJo software.

#### Fluorescence image quantification

Multiple z-stacks were imaged using Zeiss Axio Zoom.V16 with Axiocam 506 mono camera with total magnification 80x. Orthogonal projection of z-stack images was performed in ZEN Blue 2.3 software. The area of fluorescent cells in CHT or whole embryos was analyzed in Fiji using the Threshold and Analyze particles features.

#### Statistical analysis

P values were calculated by unpaired two-tailed Student's t-test using the GraphPad Prism software. Graphs depict median values or mean values with standard deviation.

### Supplemental tables

**Supplemental Table 1: Primers for cloning of *csf1r* ligands**

| | Forward primer | Reverse primer |
|---|---|---|
| **mRNA** | | |
| ***csf1a*** | CACCATGAACACACACATAACAGCCC | CCTAGGATGAAGATGCTTTAGATGATCCA |
| ***csf1b*** | GCGGATCCTTCTCCTGCTGGATCGGTGA | GCGGATCCAACACGTCTGTTTCGCTGT |
| ***il34*** | GCGGATCCACCATGGTCCAGTCCGAATG | GCGGATCCCTGCTGCTATGTTGTTTTCCT |
| **protein** | | |
| ***csf1a*** | CGGGATCCGCTGGTGTGCCAGGTCCATGTAA | CGGGATCCTATGATCCAATGCTCACTGTTG GACAAA |
| ***csf1b*** | GCGGATCCGACATCCCCGGTCCTTGCAA | GCGGATCCTTAGAAGGCTGTGGAGAGGT |
| ***il34*** | GCGGATCCGCGGCTCCAGATCTCTGTGGA | GCGGATCCCTGCTGCTATGTTGTTTTCCT |

**Supplemental Table 2: Primers for CRISPR mutant generation and genotyping**

| | | |
|---|---|---|
| **OA83** | TAGGAGTACACAGAGGATTACC | csf1a CRISPR sgRNA template |
| **OA84** | AAACGGTAATCCTCTGTGTACT | |
| **OA85** | TAGGGGATGTCCATCATGGAGA | csf1b CRISPR sgRNA template |
| **OA86** | AAACTCTCCATGATGGACATCC | |
| **OA87** | AAAATCCACGTGACTGTGCC | csf1a genotyping |
| **OA88** | CCCAAACTCACAACCTTGCA | |
| **OA89** | TGTTTGTCTTTGCGTGTCACT | csf1b genotyping |
| **OA90** | GCGTTTGAAAGGAAAAGGTGC | |
| **Tue1402** | TAGGCCTTTAACCTGGTCGGTC | csf1ra CRISPR sgRNA template |
| **Tue1403** | AAACGACCGACCAGGTTAAAGG | |
| **Tue1874** | GTTCCAGAAGGAAAGTTTC | csf1ra genotyping |
| **Tue1875** | ATATTTACTGTCATCATGGC | |

**Supplemental Table 3: Primers for qPCR**

| | Forward primer | Reverse primer |
|---|---|---|
| ***ef1a*** | GAGAAGTTCGAGAAGGAAGC | CGTAGTATTTGCTGGTCTCG |
| ***mob4*** | CACCCGTTTCGTGATGAAGTACAA | GTTAAGCAGGATTTACAATGGAG |
| ***csf1ra*** | CCATCGAGAGCATCCTGACC | ATCCACATCGTCTCCTTCGC |
| ***csf1rb*** | AGAGATTTGGCAGCACGGAA | AGAACTGGGCATCAATCGCA |
| ***mpeg1*** | CCCACCAAGTGAAAGAGG | GTGTTTGATTGTTTTCAATGG |
| ***mfap4*** | TGCTCTCAGATGGGAAAGATG | GCCAGTATTCTCCCTCCACA |
| ***l-plastin*** | AGAAGCAGCACATCAACG | AAGTGGGGTCAGAGTAATCC |
| ***mpx*** | TCAATATGAGGACGCCGTTTCT | GAATGCGATTGGAAACCAGTCT |
| ***spi1b*** | AGGAGTGTATGAGAGACCACATCAG | ATTTCGCAGAAGGTCAAGCA |

#### Supplemental figures

**A**

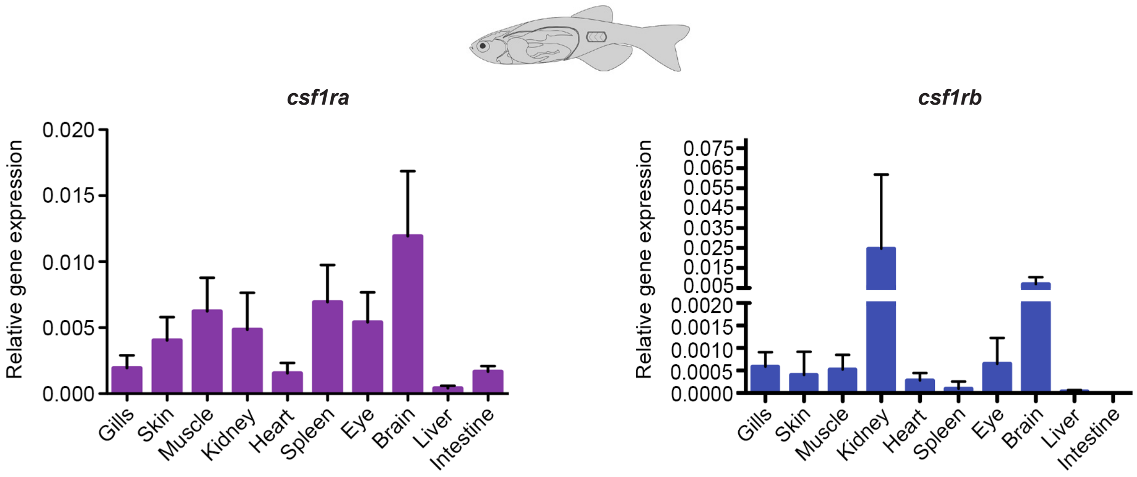


**Supplemental Figure S1: Expression of *csf1ra* and *csf1rb* in zebrafish adult tissues.**

(A) qRT-PCR analysis of *csf1ra* and *csf1rb* expression in adult zebrafish tissues. Pool of 3-5 fish organs/samples were collected in 3-5 biological replicates. The expression was normalized to *ef1α* gene.

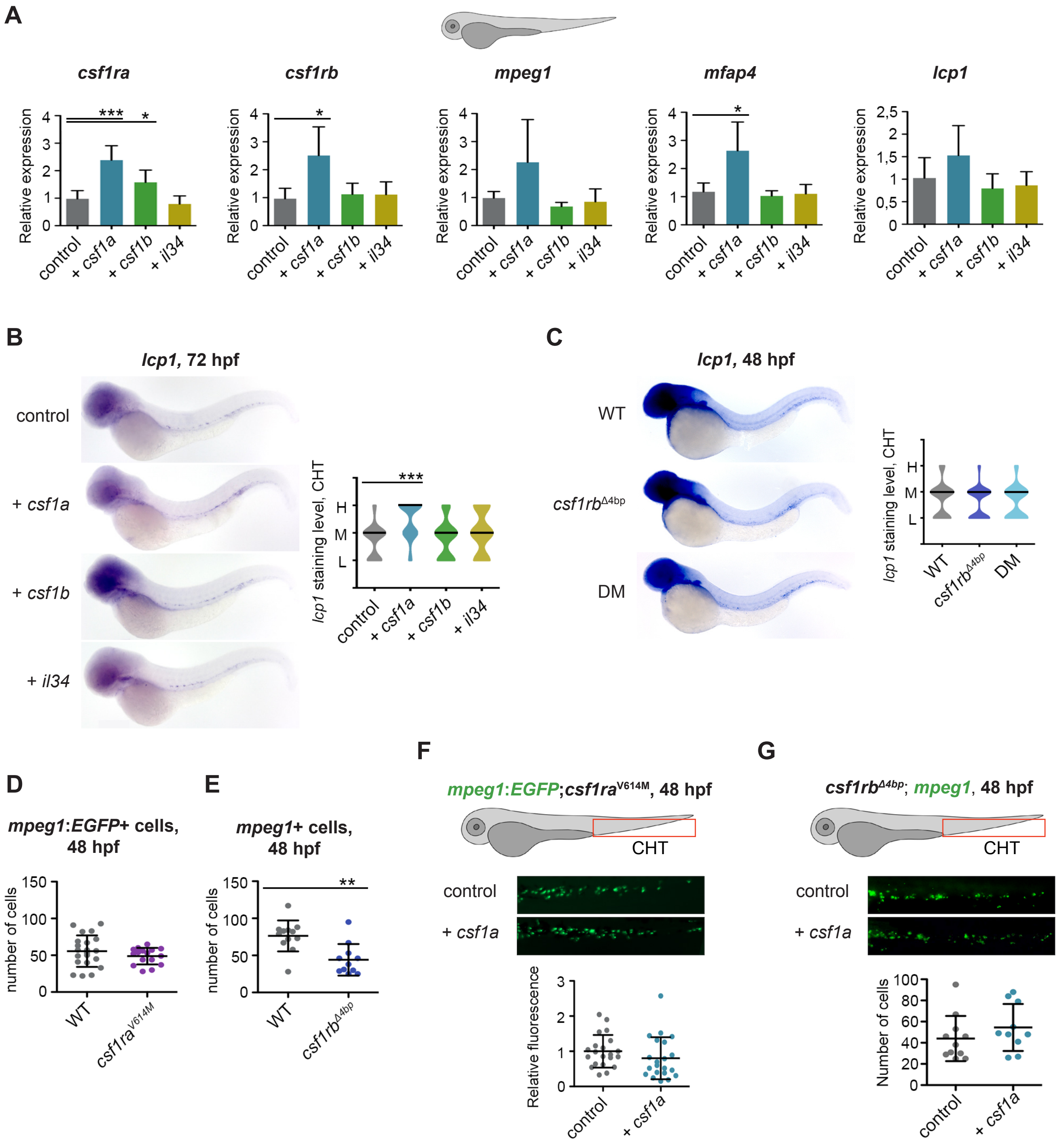


**Supplemental Figure S2: *csf1a*, but not *csf1b* or *il34* drive the expansion of embryonic macrophages.** (A-B) *csf1a, csf1b* and *il34* ligands were overexpressed by mRNA microinjection in 1-cell stage embryos. Control embryos were injected with PBS. Level of statistical significance was determined by unpaired two-tailed t-test. (A) ~20 embryos were pooled at 72 hpf and whole RNA was isolated. qRT-PCR analysis of *csf1ra, csf1rb* and macrophage specific markers - *mpeg1, mfap4* and *lcp1* was performed with 4-6 biological replicates. *P < 0.04, ***P < 0.001. (B) WISH for *lcp1* expression was performed at 72 hpf. Violin plots show the level of *lcp1* staining in the caudal hematopoietic tissue (CHT) of individual embryos (L = low, M = medium, H = high) with median represented by a black line. ***P < 0.0001. (C) WISH of 48 hpf embryos showing the expression of *lcp1* in WT, *csf1rb*$^{Δ4bp}$ and *csf1ra*$^{V614M}$;*csf1rb*$^{Δ4bp}$ double mutant (DM). Violin plots show the level of *lcp1* expression in the CHT of individual embryos (L = low, M = medium, H = high) with median represented by a black line. (D-E) Number of *mpeg1+* macrophages in the CHT of 48 hpf embryos. *mpeg1+* positive cells were counted in (D) Tg(*mpeg1:EGFP*) = WT and

Tg(*mpeg1:EGFP*);*csf1ra*$^{V614M}$ = *csf1ra*$^{V614M}$; and in (E) WT and *csf1rb*$^{\Delta 4bp}$ embryos after HCR WISH with *mpeg1* probe. **P < 0.006. (F-G) Overexpression of *csf1a* ligand by *csf1a* mRNA microinjection in 1-cell stage receptor mutant embryos. Control embryos were injected with PBS. Fluorescence images were acquired at 48 hpf and analyzed in the CHT. (F) *csf1ra* mutant transgenic embryos Tg(*mpeg1:EGFP*);*csf1ra*$^{V614M}$. The area of fluorescent cells was calculated in FIJI and the results were normalized to injected control. (G) HCR WISH of 48 hpf embryos for *mpeg1* (green) gene in *csf1rb*$^{\Delta 4bp}$ mutants. *mpeg1*+ cells in CHT were manually counted.

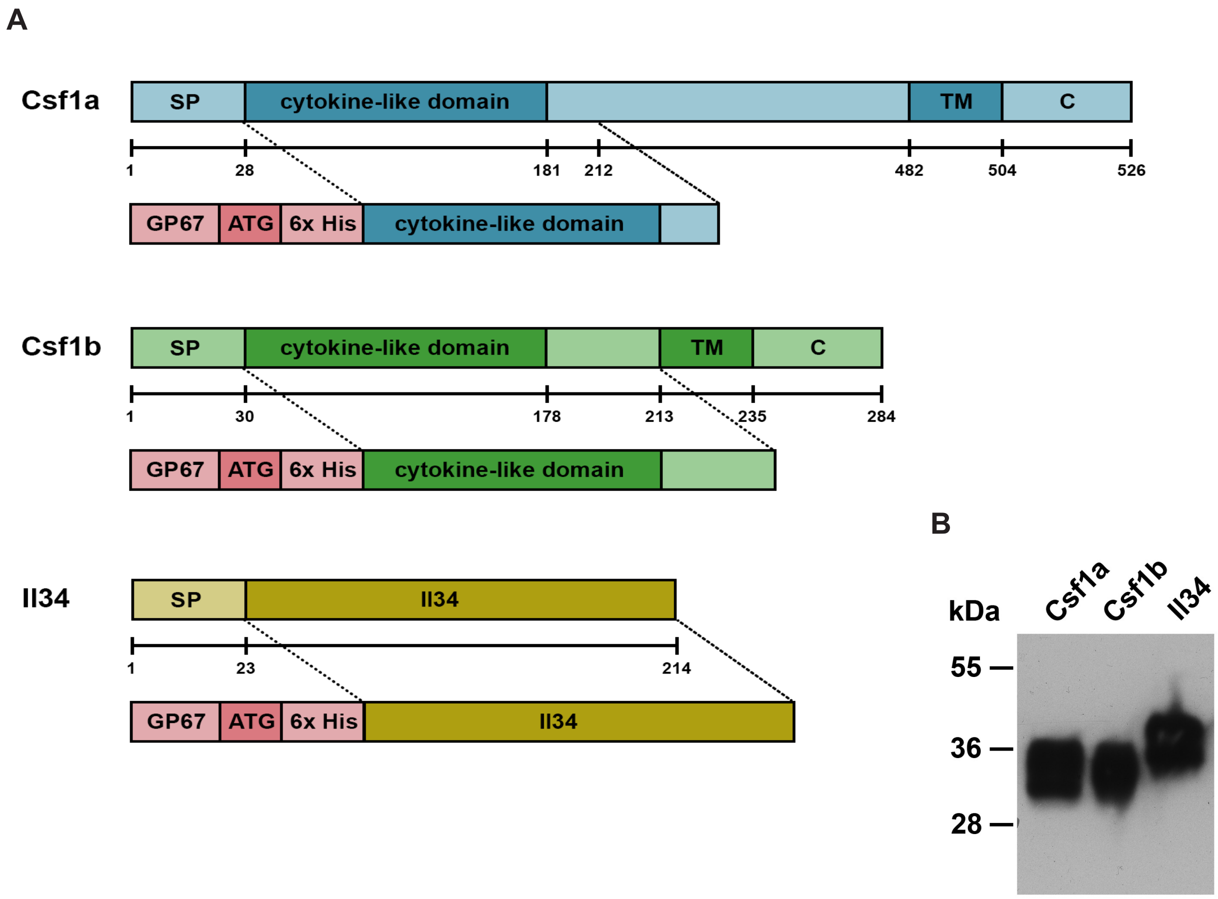


**Supplemental Figure S3: Cloning and expression of zebrafish recombinant Csf1a, Csf1b and Il34 proteins.**

(A) Cytokine-like domain without signal peptide - SP, transmembrane domain - TM and cytosolic part - C of zebrafish *csf1a* and *csf1b,* and *il34* without SP were amplified by RT-PCR from adult zebrafish organs and cloned to pAc-GP67-B vector containing 6xHis tag for expression of recombinant proteins using Baculovirus expression system in sf21 cells. (B) His-tagged recombinant Csf1a, Csf1b and Il34 proteins were purified from sf21 supernatants using affinity chromatography and detected using anti-His antibody.

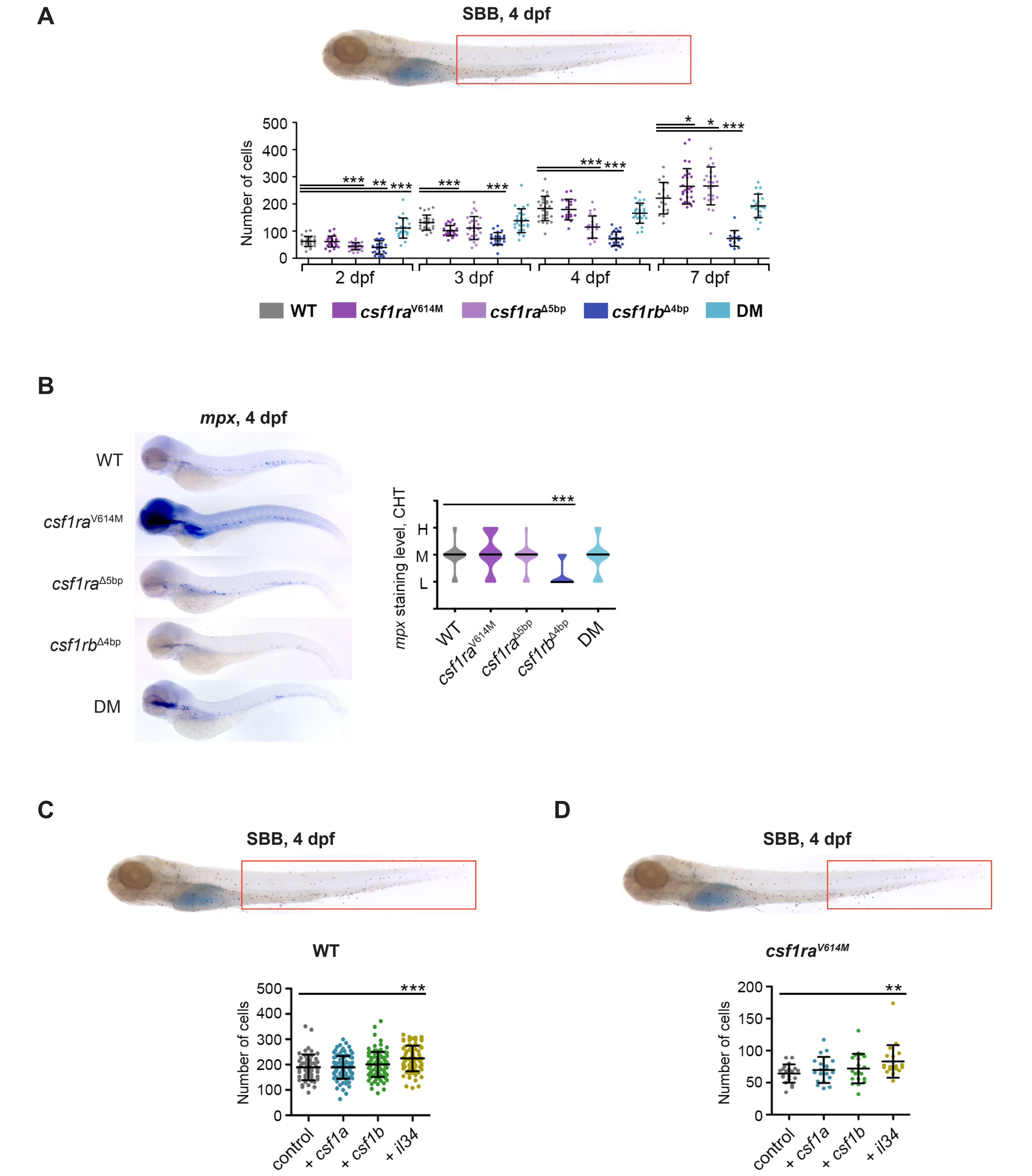


**Supplemental Figure S4: *il34* and *csf1rb* are important for embryonic neutrophil development.**

(A) Sudan black B (SBB) staining in WT, *csf1ra*$^{V614M}$, *csf1ra*$^{\Delta 5bp}$, *csf1rb*$^{\Delta 4bp}$ and *csf1ra*$^{V614M}$;*csf1rb*$^{\Delta 4bp}$ (DM) 2-7 dpf embryos. The analyzed area of the larvae is marked with a red rectangle. The graph shows the number of SBB positive cells. SBB positive cells were manually counted, and the level of statistical significance was determined by unpaired two-tailed t-test. *P < 0.04, **P < 0.006, ***P < 0.0001. (B) WISH of 4 dpf embryos showing the expression of *mpx* in WT, *csf1ra*$^{V614M}$, *csf1ra*$^{\Delta 5bp}$, *csf1rb*$^{\Delta 4bp}$ and *csf1ra*$^{V614M}$;*csf1rb*$^{\Delta 4bp}$ (DM) embryos. Violin plots show the level of *mpx* expression in individual embryos (L = low, M = medium, H = high) with median represented by a black line. ***P < 0.0001. (C-D) SBB staining in 4 dpf embryos. The analyzed area is marked with a red rectangle. SBB positive cells were manually counted, and the level of statistical significance was determined by unpaired two-tailed t-test. (C) *csf1a, csf1b* and *il34* ligands were overexpressed by mRNA microinjection in 1-cell stage WT embryos. Control embryos were injected with PBS. The graph shows the number of SBB positive cells. ***P < 0.001. (D) *csf1a, csf1b* and *il34* ligands were overexpressed by mRNA microinjection in 1-cell stage *csf1ra*$^{V614M}$ mutant embryos. Control embryos were injected with PBS. The graph shows the number of SBB positive cells. **P < 0,006. All SBB staining images were acquired on Zeiss Axio Zoom.V16 with Zeiss Axiocam 105 color camera and processed using the Extended Depth of Focus module in the ZEN blue software. FIJI and Adobe Photoshop were used for image processing.

**A**

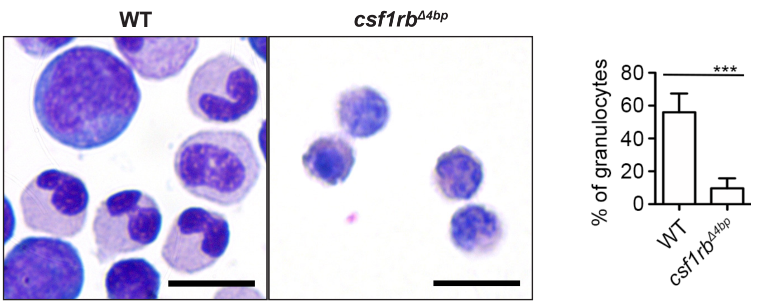


**Supplemental Figure S5: *csf1rb* is required for proper heterophil maturation.**
(A) Smears of WKM cells from WT and *csf1rb*$^{\Delta 4bp}$ fish were stained with May-Grünwald and Giemsa. 100 cells were counted in 3-5 biological replicates. The graph shows the percentage of lobulated heterophils. The scale bar on the microscopic images is 10 µm. Bright field images were acquired on Leica DM 2000 microscope with Zeiss Axiocam 105 color camera.

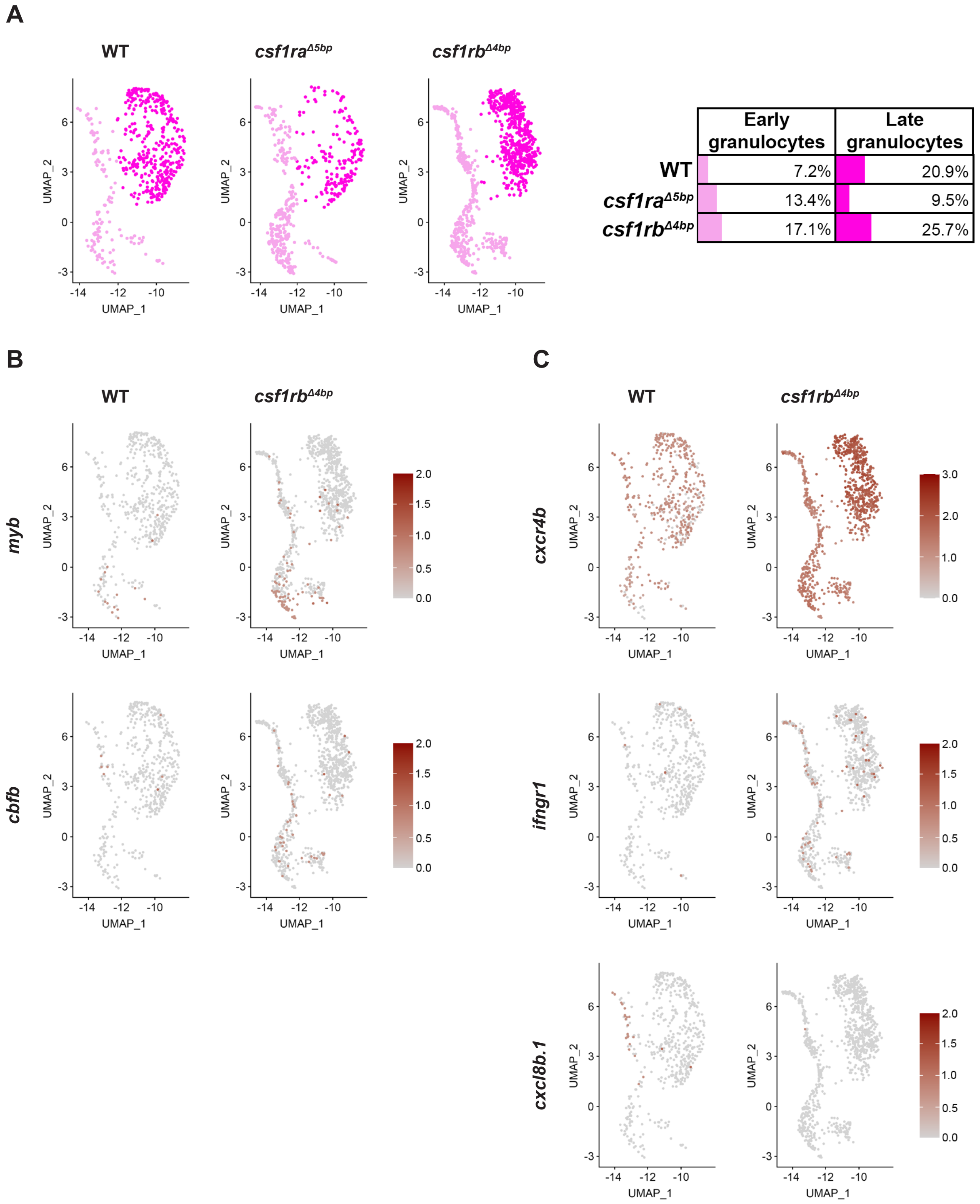


**Supplemental Figure S6: The maturation and migration of granulocytic cells is impaired in *csf1rb*$^{\Delta 4bp}$**

(A) Clusters in Dim plots represent early-like (light pink) and late-like (magenta) granulocyte populations of WT, *csf1ra*$^{\Delta 5bp}$ and *csf1rb*$^{\Delta 4bp}$ fish. The percentage of cells in each subpopulation is displayed in the table. (B-C) Feature plots of the granulocyte population in WT and *csf1rb*$^{\Delta 4bp}$ fish show the expression level of selected marker genes from gray (lowest) to red (highest). (B) Progenitor/early-like markers. (C) Migration and inflammation markers.

### References

1. Stachura DL, Svoboda O, Lau RP, et al. Clonal analysis of hematopoietic progenitor cells in the zebrafish. *Blood*. 2011;118(5):1274-1282.
2. Svoboda O, Stachura DL, Machonova O, Zon LI, Traver D, Bartunek P. Ex vivo tools for the clonal analysis of zebrafish hematopoiesis. *Nat Protoc*. 2016;11(5):1007-1020.
3. Svoboda O, Stachura DL, Machonova O, et al. Dissection of vertebrate hematopoiesis using zebrafish thrombopoietin. *Blood*. 2014;124(2):220-228.
4. Irion U, Krauss J, Nusslein-Volhard C. Precise and efficient genome editing in zebrafish using the CRISPR/Cas9 system. *Development*. 2014;141(24):4827-4830.
5. Thisse C, Thisse B. High-resolution in situ hybridization to whole-mount zebrafish embryos. *Nat Protoc*. 2008;3(1):59-69.
6. Kafina MD, Paw BH. Using the Zebrafish as an Approach to Examine the Mechanisms of Vertebrate Erythropoiesis. *Methods Mol Biol*. 2018;1698:11-36.
7. Le Guyader D, Redd MJ, Colucci-Guyon E, et al. Origins and unconventional behavior of neutrophils in developing zebrafish. *Blood*. 2008;111(1):132-141.
